# Angiopoietin-like protein 7 (ANGPTL7)-mediated effects on resident immune cells contribute to neurodegeneration in glaucomatous optic nerve head

**DOI:** 10.64898/2026.09.27.751872

**Authors:** Thomas R. Vollmer, Katie Dolan, Chuanxi Xiang, Angela Banks, Eileen M. Cooney, Jessica Heyder, Charles C. King, Joanna Vrouvlianis, Akshata N. Gujar, Nathaniel D. Kirkpatrick, Xiaoqiu Wu, Carter Jones, Loren J. Miraglia, Anthony P. Orth, Yubin Qiu, Melissa Prentiss, Isra S. Hassan, Julian Kwan, Roxana Garcia Caro, Lin Fan, Yanqun Wang, Sheng-Yong Niu, Chia-Ling Huang, VijayKrishna Raghunathan, Amy Chen, Nhi Vo, Angela Chui, Robert Esterberg, Ganesh Prasanna, Christopher W. Wilson

## Abstract

Glaucoma is an optic neuropathy characterized by progressive retinal degeneration. Population genetic analyses describe an association of variants in angiopoietin-like protein 7 (ANGPTL7) with lower intraocular pressure (IOP) and decreased glaucoma susceptibility. In addition to its reported role in the trabecular meshwork (TM), we demonstrate that ANGPTL7 is also expressed within disease-relevant cell types of the human posterior eye, specifically prelaminar optic nerve head astrocytes (ONHA). Contrasting with previously published findings, we observe no difference in IOP between *Angptl7^-/-^* mice and wild-type littermates. Despite absence of an IOP phenotype, loss of ANGPTL7 protects from IOP-induced retinal ganglion cell (RGC) damage in a rodent glaucoma neurodegeneration model. ANGPTL7 may signal through LAIR1-positive immune cells, potentially contributing to pro-inflammatory signaling and phenotypic modulation of resident immune cells, with evidence suggesting that oligomerization of ANGPTL7 is necessary for its activity. Comprehensively, our data highlight an unappreciated role of ANGPTL7 in modulating neuroinflammation in the posterior eye, contributing to glaucoma progression.

## Introduction

Glaucoma is a degenerative optic neuropathy characterized by retinal ganglion cell (RGC) loss and cupping of the optic nerve head (ONH), presenting clinically with peripheral vision loss followed by progressive restriction in the central visual field ultimately leading to blindness [1]. Glial cells (astrocytes, microglia) within the neural retina and optic nerve tissue can become activated in glaucoma, initiating a neuroinflammatory cascade of cytokine/chemokine production and increased phagocytosis and fibrosis, creating a neurotoxic environment that can ultimately damage RGCs [2]. The most prevalent form of glaucoma is primary open-angle glaucoma (POAG), which is associated with elevated intraocular pressure (IOP). IOP is the only modifiable risk factor as presently there are no approved therapies that directly protect RGCs and the optic nerve. IOP-lowering drugs offer a palliative, non-curative option for most POAG patients, however their effectiveness can suffer from poor patient compliance with daily eye drops and disease progression despite IOP-lowering [3-5]. These situations highlight an overarching unmet medical need to identify therapies directly targeting neuroinflammatory milieu as a way of treating disease independent of IOP-lowering therapies.

Multiple population studies have identified protein-altering genetic variants in angiopoietin-like protein 7 (ANGPTL7) that are associated with lower IOP and reduced incidence of glaucoma [6, 7]. Mechanistic studies suggest that these are loss-of-function variants that destabilize and/or result in decreased secretion of ANGPTL7 [7]. ANGPTL7 is increased in aqueous humor of POAG patients and in glaucomatous preclinical models [8], and levels of *ANGPTL7* are increased in patient-derived trabecular meshwork cells *in vitro* and donor explants following glucocorticoid treatment [9, 10]. Preclinical assessments of IOP in *Angptl7^-/-^*mice have yielded conflicting results [11, 12], with increased or decreased IOP observed in knockouts. However, ANGPTL7-IOP association does not necessarily preclude an alternative site of action elsewhere in the eye. For example, ANGPTL7 transcript and protein levels were detected and elevated in the cribriform plates of the lamina cribrosa at the ONH in sections from a donor with IOP-associated glaucoma secondary to neovascularization [8], suggesting a possible role within the posterior eye. A clearer mechanistic understanding of ANGPTL7 is thus critical towards establishing therapeutic hypotheses surrounding this potential glaucoma target, which may include both an IOP-dependent (anterior eye) and a putative IOP-independent role in neuroinflammation (posterior eye).

ANGPTL7 is a secreted glycoprotein [8, 13], though the nature of interacting partners and downstream effector functions remains elusive. The hypothesis that glaucoma-protective ANGPTL7 variants are not secreted [6, 7] suggests that interaction with an extracellular factor likely mediates normal ANGPTL7 function within the eye. Studies in mice point to a requirement of ANGPTL7 for signaling to cells of the myeloid lineage [14], and are bolstered by *in vitro* findings indicating binding of ANGPTL7 to the leukocyte receptors LAIR1 (leukocyte-associated immunoglobulin-like receptor 1) and LILRB2 (leukocyte immunoglobulin-like receptor subfamily B member 2) when exogenously expressed [15]. The ONH region is a crucial site for crosstalk between glial cells [16-18] and these links to immune cells suggest a possible role for ANGPTL7 signaling to glia in neuroinflammatory glaucoma.

Here, we describe a novel role for ANGPTL7 in promoting proinflammatory cytokine release from cells of immune origin, which may occur in the ONH region to drive neuroinflammation via astroglial activation and RGC loss. Using a combination of single nuclear RNA sequencing (snRNAseq) and *in situ* hybridization in human donor eyes, we show that *ANGPTL7* transcript is expressed in a subset of human ONH astrocytes (ONHA) and is present in glaucomatous tissue. In an ocular hypertension model of glaucoma neurodegeneration [19], genetic deletion of ANGPTL7 protects from IOP-induced RGC loss and gliosis. Mechanistically, oligomerized ANGPTL7 stimulates pro-inflammatory signaling and promotes adhesion in an immune cell line. Taken together, our data support a direct role for ANGPTL7 in mediating neuroinflammatory processes and suggest that inhibition of ANGPTL7 at the ONH may provide a direct neuroprotective therapeutic option.

## Methods

### Human donor eye acquisition and histology

Deidentified human donor eyes were obtained from the Lions World Vision Institute (Tampa, FL) with informed consent of donors or donors’ next of kin, in accordance with the Eye Bank Association of America (EBAA) medical standards, US/Florida law for human tissue donation, the Declaration of Helsinki and Food and Drug Administration (FDA) regulations, and guidance from the Novartis Biosample compliance team. Whole globes were enucleated <6.5 hours postmortem and were evaluated for not meeting criteria for corneal transplantation. Globes were fixed at room temperature with Modified Davidson’s Fixative (MDF) [20] for 48 hours and were shipped to Novartis in MDF, followed by 70% ethanol incubation for an additional 48 hours. Afterwards, eyes were dehydrated and embedded in paraffin with a Tissue-Tek VIP and Tissue-Tek Tec (Sakura Finetek, Torrance, CA). Paraffin sections (5 µm) were cut, processed and graded (where applicable) for glaucoma severity using a multiparametric approach [20]. Fresh sections were then obtained and used for RNAscope.

RNAscope probes are detailed in Supplemental Table 8. RNAscope staining for all probes was carried out using an Automated Ventana immunohistochemistry (IHC) staining instrument (Roche Diagnostics, Indianapolis, IN) according to manufacturer’s instruction (Advanced Cell Diagnostics (ACD), Newark, CA). RNA signal was detected by RNAscope VS Universal AP Reagent kit (ACD, Newark, CA) and mRNA Red Detection kit (Roche Diagnostics, Indianapolis, IN). Brightfield images for chromogenic slides were collected via Aperio A2 scanner (Leica, Wetzlar, Germany). For co-staining data, immunofluorescent (IF) staining with IBA1 antibody was run after completion of RNAscope. IF signal was revealed via OmniMap and Discovery FAM kits (Roche Diagnostics, Indianapolis, IN). Hoechst (Thermo Scientific, Waltham, MA) was used for nuclear staining of immunofluorescence slides. Fluorescent slides were scanned and imaged using an AxioScan Z1 (Zeiss, Oberkochen, Germany).

### Human donor tissue acquisition, dissection and single-nuclei RNA-sequencing (snRNAseq)

Additional human ocular tissues were procured as described above and dissected by the Lions World Vision Institute (Tampa, FL). Briefly, a 6 mm biopsy punch centered on the fovea was used to collect macular neural retina tissue. Optic nerve was cut either flush with or 1 mm from sclera and ONH was collected via biopsy punch. Tissues were flash-frozen, stored, and shipped to Novartis at -80 °C. Tissues were thawed and nuclei were extracted using a modified version of the ‘Frankenstein’ protocol [21]. Optic nerve and ONH tissues were subjected to flow cytometry to remove debris and to enrich nuclei prior to barcoding. snRNAseq libraries were constructed with 3’ Gene Expression kits (v3) from 10X Genomics (Pleasanton, CA) according to manufacturer’s instructions. Libraries were QC’d using a High Sensitivity DNA kit on a Bioanalyzer (Agilent Technologies, Lexington, MA). Libraries were pooled and sequenced on a NovaSeq (Illumina, San Diego, CA) per manufacturer’s instructions. Data were processed using 10X Genomics CellRanger (Version 5.0.1 and 6.0.2), exons and introns were mapped to human reference genome (GRCh38). Downstream analyses were carried out using Seurat (Version 4.3.0) in the R environment (R Version 4.1.0).

### RNAscope, immunohistochemistry (IHC), and image analysis

Arivis Vision 4D 4.1.1 (Zeiss, Oberkochen, Germany) software was used to visualize and compare stained regions for *ANGPTL7* (RNAscope), *GFAP* (RNAscope), and IBA1 (immunofluorescence) expression patterns in the ONH. A custom analysis pipeline was developed to 1) segment the cell nuclei based on Hoechst stain, 2) segment the markers of interest (*ANGPTL7*, *GFAP* & IBA1) through the object detector tool, 3) identify *GFAP*- and IBA1-positive objects (cells), and 4) measure co-localization of *ANGPTL7*-positive spots within the identified cell subsets; *GFAP*-positive cells were derived from RNAscope spots, so the pipeline used a local smoothing and region growing feature to fill in the cell region of *GFAP* positivity. All data presented are as a percentage of *ANGPTL7*-positive spots within *GFAP*-positive or IBA1-positive objects. Representative images of co-staining and quantification are provided in Supplemental Fig. 3c and 3d, respectively.

### Establishment of *Angptl7^-/-^* mice and wild-type littermates

All rodent studies were conducted in conformity with Institutional Animal Care and Use Committee (IACUC) guidelines. *Angptl7^-/-^* mice were originally generated by Lexicon Pharmaceuticals, Inc. (The Woodlands, Texas) on a mixed background of C57BL/6J X 129S5 mice via gene targeting by homologous recombination in exon 1 [22]. *Angptl7^-/-^* mice were cryo-recovered from sperm on a mixed 129S5 x C57BL/6J-Tyrc-Brd background by Taconic Biosciences (Germantown, NY). Female oocyte donors were subjected to *in vitro* fertilization (IVF) with frozen sperm of *Angptl7^-/-^* mice, after which heterozygous offspring were crossed to generate homozygous knockouts (*Angptl7^-/-^*), heterozygous knockouts (*Angptl7^+/-^*) and wild-type littermate controls. Genotypes were confirmed upon delivery by conventional genotyping, quantitative PCR (qPCR) and RNAscope (Supplemental Fig. 4).

### Intraocular pressure (IOP) measurement

IOP was measured in mice under anesthesia in the morning (between 7 a.m. and noon) using a rebound Tonometer (TonoLab, Tiolat, Oy, Finland) calibrated for measurement on murine eyes. Briefly, animals were anesthetized with 3% isoflurane supplemented with oxygen at a flow rate of 1 liter per minute for 8 minutes. A nose cone was applied on the animals with isoflurane flowing, and IOP was measured. IOP measurements from each session are expressed as the mean of 10 successive measurements for each eye ± standard error of the mean (SEM).

### Hyaluronic acid glycidyl methacrylate (HAMA) neuroprotection model

HAMA studies were carried out as previously described [19, 23], including synthesis of HAMA, preparation of injection setup and intracameral (IC) injection followed by *in vivo* photopolymerization. HAMA solution was protected from bright light in a dark container. Sterile PBS was added to the stock HAMA solution to make a 2% working concentration. Sterile fluorescein (Akorn, Lake Forrest, IL) was added to the injection solution at a final concentration of 0.05% (w/v) to aid in visualization during injection. IOP was measured at baseline, then 3, 7, 14, 21 and 28 days following HAMA injection.

Cumulative IOP profiles for each eye were calculated. Animals with adverse clinical observations (i.e. hypoxia, neovascularization, etc.) were excluded (<10%) from further analyses, as were (1) naïve/PBS-injected animals with cumulative IOP >30% above naïve/PBS average and/or (2) HAMA-injected animals with cumulative IOP <30% above naïve/PBS average. Following completion of the 28-day study, animals were euthanized and eyes were collected for retinal flat-mount staining. Percentage RGC loss or astrocyte or microglia activation was calculated by normalizing naïve samples to 100: ((Naive – HAMA) ÷ Naïve) x 100 for each respective genotype. The delta between HAMA and naïve is described as percent RGC loss or percent astrocyte or microglia activation; percent protection is characterized as the delta between naïve and HAMA for each respective readout, i.e. WT RGC percent loss – Angptl7^-/-^ RGC percent loss = percent protection.

### Optical coherence tomography (OCT) of mouse cornea

Optical coherence tomography (OCT) images were obtained using a Bioptigen spectral-domain ophthalmic imaging system (Leica, Wetzlar, Germany). Images were acquired from *Angptl7^-/-^*and wild-type littermate mice at 3 and 5 months of age. Animals were anesthetized with intraperitoneal (IP) injection of ketamine/xylazine (100mg/kg / 5mg/kg) cocktail prior to the OCT procedure and given 1.0% cyclopentolate (Alcon, Fort Worth, TX) and 2.5% – 10% phenylephrine (Akorn, Lake Forrest, IL) topically for dilation in addition to 0.5% proparacaine (Akorn, Lake Forrest, IL) as a topical anesthetic. Animals were retained on a heating pad at sternal recumbency during the procedure and their eyes were topically hydrated with mild to moderate GenTeal (Alcon, Fort Worth, TX). Ten linear b-scans were acquired and later averaged. One horizontal (nasal to temporal) and one vertical (inferior to superior) linear b-scan series was acquired in a cross pattern centered on the central cornea. Morphometric analysis of OCT images was performed on horizontal b-scans using a custom MATLAB algorithm (Mathworks, Natick, MA – Version R2022a) to align and average b-scans, after which, user input highlighted the layers to be quantified for thickness.

### Preparation of retinal flat-mounts for immunofluorescent staining (mouse tissue)

After euthanasia, eyes were enucleated and retinal flat-mounts were processed as previously described [19, 23]. Whole globes were fixed in 4% paraformaldehyde (PFA) for two hours, after which neural retina was dissected out and cut into four petals to lie flat. Each flat-mount was stained for BRN3A (retinal ganglion cell marker), GFAP (astrocyte marker), and IBA1 (microglia marker). Antibodies used for staining are described in Supplemental Tables 5 and 6. Primary antibody dilutions were prepared in eBioscience IHC/ICC blocking buffer (Invitrogen, Waltham, MA) and incubated for 48 hours at 4 °C. Secondary antibody incubations were prepared in PBS and incubated overnight at 4 °C. Fluorescent images of flat-mounts were obtained using an Axioscan Z1 (Zeiss, Oberkochen, Germany) at 10X magnification. Images were stitched together to show the whole retina for each respective marker.

### Quantification of retinal flat-mount staining

Prior to quantification, flat-mount images were visually inspected using ZEN software (Zeiss, Oberkochen, Germany) to ensure background uniformity, picture clarity and focus. Custom CellProfiler [24] (Broad Institute, Cambridge, MA) pipelines were designed for each respective readout. Retinal degeneration was measured via BRN3A staining of RGC nuclei; nuclei were quantified and reported as RGC density (RGC/mm^2^). GFAP staining was used to measure astrocyte coverage area and is reported as astrocyte area (%). Microglia were stained with IBA1; quantification algorithm counts the number of IBA1-positive cells per unit area and is reported as microglia area (Microglia/mm^2^).

### Cell culture

THP-1 human monocytes and U-2 OS human bone osteosarcoma cells were obtained from American Type Culture Collection (ATCC, Manassas, VA). THP-1 cells were cultured in RPMI-1640 (ATCC) supplemented with 10% heat-inactivated fetal bovine serum (FBS), 1% MEM non-essential amino acids, 1% penicillin-streptomycin-glutamine (PSG), 1% sodium pyruvate and 0.1 mM beta-mercaptoethanol (Thermo Fisher Scientific, Waltham, MA). For immunofluorescent binding experiments, THP-1 cells were differentiated for 24 to 48 hours with 100 nM phorbol myristate acetate (PMA) (Millipore-Sigma, Burlington, MA). U-2 OS cells were cultured in McCoy’s 5A media (Invitrogen, Waltham, MA) supplemented with 10% FBS and 1% PSG.

### ANGPTL7 *in vitro* binding experiments

ANGPTL7 protein was generated internally according to Gonzales et al [25]. The ANGPTL7 gene was cloned into a vector with an N-terminal or C-terminal mouse immunoglobulin Fc domain, described as N-Fc ANGPTL7 and ANGPTL7 C-Fc, respectively. Angiopoietin 2 (ANG2) gene was cloned with an N-terminal tag only (N-Fc ANG2). Protein expression and size was validated via Western blot (Supplemental Fig. 9). An anti-mouse Fc FITC-conjugated antibody (Millipore-Sigma, Burlington, MA) was validated for detection of Fc tag via immunofluorescence following overexpression in U-2 OS cells (Supplemental Fig. 10). To facilitate ectopic receptor over-expression evaluation, canonical open-reading frame sequences for LILRA2, LILRA3, LILRA4, LILRA5, LILRA6, LILRB2, LAIR1, and TEK were synthesized, cloned without tags into pDEST40 backbone, sequence confirmed, prepared as endotoxin-free plasmid DNA and transiently transfected into U-2 OS cells, after which N-Fc ANGPTL7 was added and incubated for 1 hour. THP-1 cells were used as a LAIR1/LILRB2 endogenously-expressing line; PMA-treated for immunofluorescence, naïve for FACS. For immunofluorescence-based imaging experiments, cells were fixed for 10 minutes with 4% PFA before blocking in tris-buffered saline (TBS) with 10% goat serum (Invitrogen, Waltham MA) for 1 hour at room temperature. Cells were incubated with an anti-mouse Fc FITC-conjugated antibody overnight at 4 °C after which 20X fluorescent images were collected using an InCell2200 (Cytivia, Washington D.C.). For FACS studies, cells were incubated with Purified NA/LE Human BD Fc block (Becton Dickinson, Waltham, MA), followed by addition of ANGPTL7 or control proteins (1 hour incubation), after which cells were incubated with FITC-conjugated anti-mouse Fc (Sigma Aldrich, St. Louis, MO) for 1 hour. All steps were carried out on ice in PBS supplemented with 0.1% BSA. FACS analysis was carried out using FlowJo software (FlowJo LLC, Ashland, OR). Cells were gated using forward and side-scatter to filter debris, then gated to filter out doublet cells.

### THP-1 mechanistic studies with oligomerized ANGPTL7

For mechanistic studies, THP-1 cells were treated with a dilution series (300, 900 or 2,700 ng/mL) of recombinant human ANGPTL7 (rhANGPTL7) (R&D Systems, Minneapolis, MN) (6X His-tagged) with or without oligomerization. Cross-linkage/oligomerization of rhANGPTL7 was achieved by pre-incubating ligand with an anti-6X His antibody before administering to cells; rhANGPTL7 (300, 900 or 2,700 ng/mL) was incubated with 5 µg/mL anti-His antibody in complete RPMI media for 5 minutes before adding to cells; non-oligomerized rhANGPTL7 was prepared in parallel without His antibody. For transcript measurement via qPCR, samples were treated for 48 hours in 6-well plates at a density of 350,000 cells per well. Plates were scraped to collect suspension cells and cells that attached post-treatment. RNA was extracted using a Direct-zol kit (Zymo Research, Irvine, CA). RNA was quantified using a microvolume spectrophotometer (NanoDrop, ThermoFisher Scientific, Waltham, MA). An equivalent amount of RNA was used to construct cDNA libraries with SuperScript IV VILO MasterMix (ThermoFisher Scientific, Waltham, MA), after which qPCR was run on a ViiA-7 device with TaqMan reagents (Thermo Fisher Scientific, Waltham, MA). Each well had an internal housekeeping control; experimental genes used FAM dye and housekeeping gene (*GAPDH*) used VIC dye. Experimental cycle threshold (CT) values were subtracted from housekeeping CT values to calculate ΔCT. ΔCT values were then subtracted from control ΔCT to calculate ΔΔCT. qPCR data are expressed at 2^-ΔΔCT^. Amplification curves were visually inspected to QC reported CT values; samples with CT values above 35 were flagged and filtered from further analysis. For Luminex and attachment assays, THP-1 cells were plated at a density of 70,000 cells per well in a 96-well plate and treated for 72 and 96 hours. At the time of collection, media containing suspension cells was pipette-mixed, transferred to a U-bottom plate and centrifuged for 3 minutes at 300 x g (4 °C). Conditioned media was transferred to a fresh 96-well plate and flash-frozen on dry-ice and subsequently assayed for cytokine secretion via Luminex assay (Luminex, Austin, TX). Pre-mixed cytokine panel kits were obtained from R&D Systems (Minneapolis, MN) and carried out according to manufacturer’s instructions; analytes described here were all within range of standard curves. For attachment assay, after removal of conditioned media for Luminex assay, plates were washed twice with PBS following incubation with CTG 2.0 reagent (Promega, Madison, WI) for 5 minutes before transfer to solid-white plates for luminescence measurement with an Envision plate-reader (Perkin-Elmer, Waltham, MA). Data were control-normalized with His-antibody control treatment set to 100.

### Statistics

Statistical analyses were performed using GraphPad Prism (v10.1.2) (Dotmatics, Boston, MA). Neuroprotection data (RGC, microglia and astrocyte quantification) and THP-1 functional data (qPCR, Luminex and attachment assay) used one-way ANOVA with Tukey’s multiple comparisons test. All other data used unpaired t-tests.

## Results

### Transcriptional profiling of glaucomatous human donor tissue indicates a potential role for ANGPTL7 in the posterior eye

Previous studies detected *ANGPTL7* transcript and protein in human and rodent ocular anterior chambers, specifically in scleral tissue, corneal stroma and trabecular meshwork (TM) [8, 11, 26, 27]. To validate and expand on published findings, we analyzed internally generated snRNAseq datasets using post-mortem, non-diseased human donor tissue from the anterior (limbal ring) and posterior eye (neural retina, ONH and optic nerve) (Fig. 1) [manuscripts in preparation; data not shown].

**Fig. 1.**
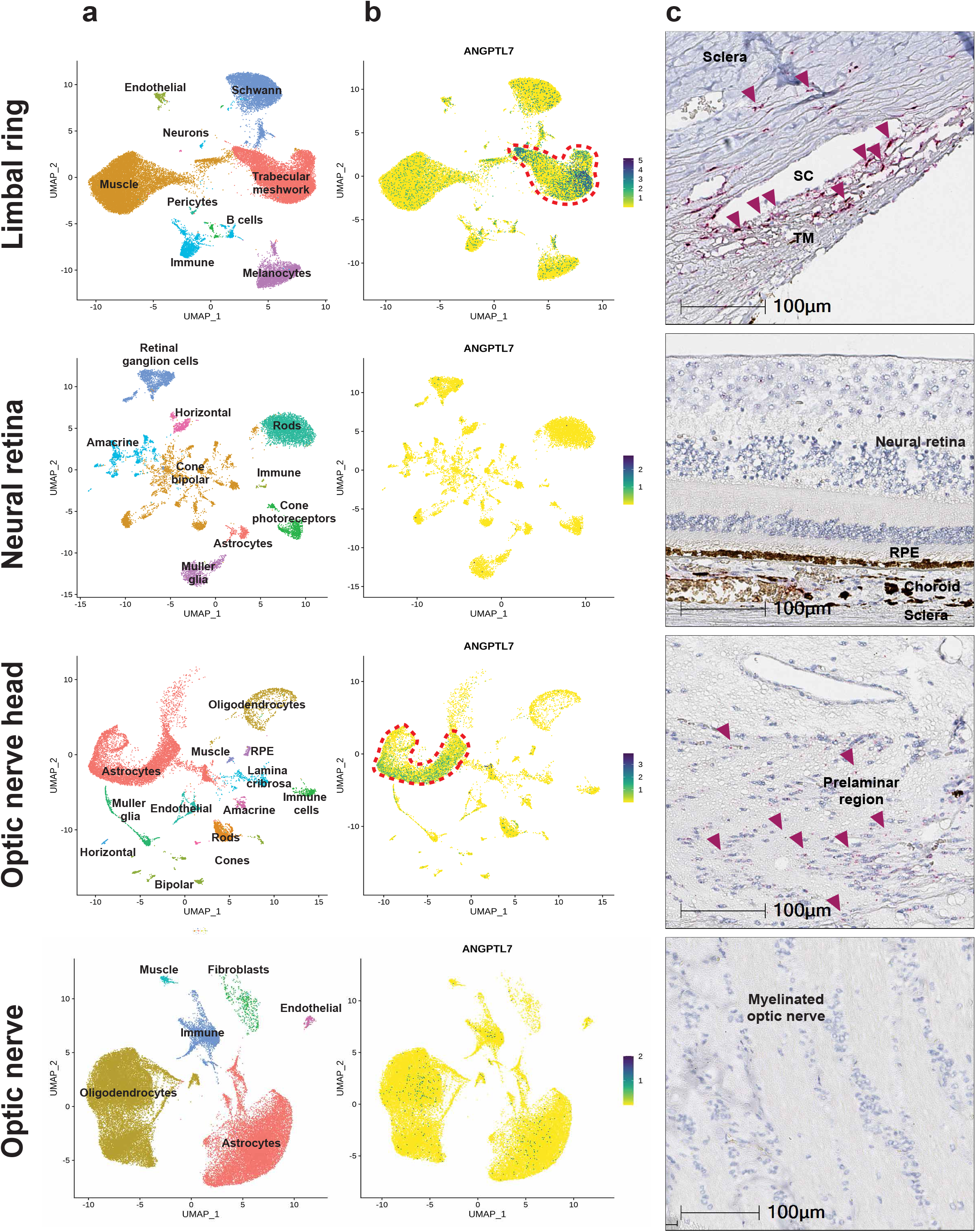
*ANGPTL7* transcript is enriched in TM cells and ONHA. snRNAseq was carried out using non-diseased post-mortem human donor material from various ocular tissues; (1) limbal ring (n = 5 donors), (2) macular neural retina (n = 2 donors), (3) ONH (n = 3 donors), (4) optic nerve (n = 3 donors); detailed donor information can be found in Supplemental Table 1. Locations from where each tissue was dissected is detailed in Supplemental Fig. 1. a UMAP depicting snRNAseq clustering of cell types within each tissue. b UMAP showing ANGPTL7 transcript levels captured from each cell type; number of transcripts per nuclei. Positive nuclei are defined as having >1 transcript per cell. c ANGPTL7 transcript levels from a non-diseased human donor (RNAscope); purple arrows highlight copies of ANGPTL7 transcript. DapB was used as a negative control stain (Supplemental Fig. 2). TM – trabecular meshwork; SC – Schlemm’s canal, RPE – retinal pigmented epithelium.

In human limbal ring tissue, *ANGPTL7* expression was predominantly found in TM cells (Fig. 1b), consistent with previously published data [26, 27]. RNAscope confirmed these findings, highlighting a concentration of transcript in the TM region with expression in the sclera and corneal stroma (Fig. 1c). In the posterior eye, 14% of ONHAs expressed *ANGPTL7*, whereas only ∼0.3% of astrocytes in the macular neural retina and 0.6% of the optic nerve astrocytes (ONA; as observed in the myelinated optic nerve) were *ANGPTL7* positive. Transcript detection via RNAscope was exclusive to the TM, prelaminar ONH region and sclera, with little to no staining in the neural retina, retinal pigmented epithelium (RPE) or myelinated regions of the optic nerve, consistent with our snRNAseq findings (Fig. 1c).

We next measured *ANGPTL7* transcript in anterior (TM) and posterior (ONH) eye of normal and POAG post-mortem human tissue sections via RNAscope to assess changes in disease. *ANGPTL7* transcript was detectable in the TM region (representative images shown in Fig. 2a, additional donors in Supplemental Fig. 3a) and prelaminar ONH (Fig. 2b, additional donors in Supplemental Fig. 3b) of both non-diseased and glaucomatous tissue. While *ANGPTL7* levels in the TM appeared similar between glaucomatous and non-diseased tissue, we noted a qualitative elevation in the prelaminar ONH of glaucomatous tissue (Fig. 2b).

**Fig. 2.**
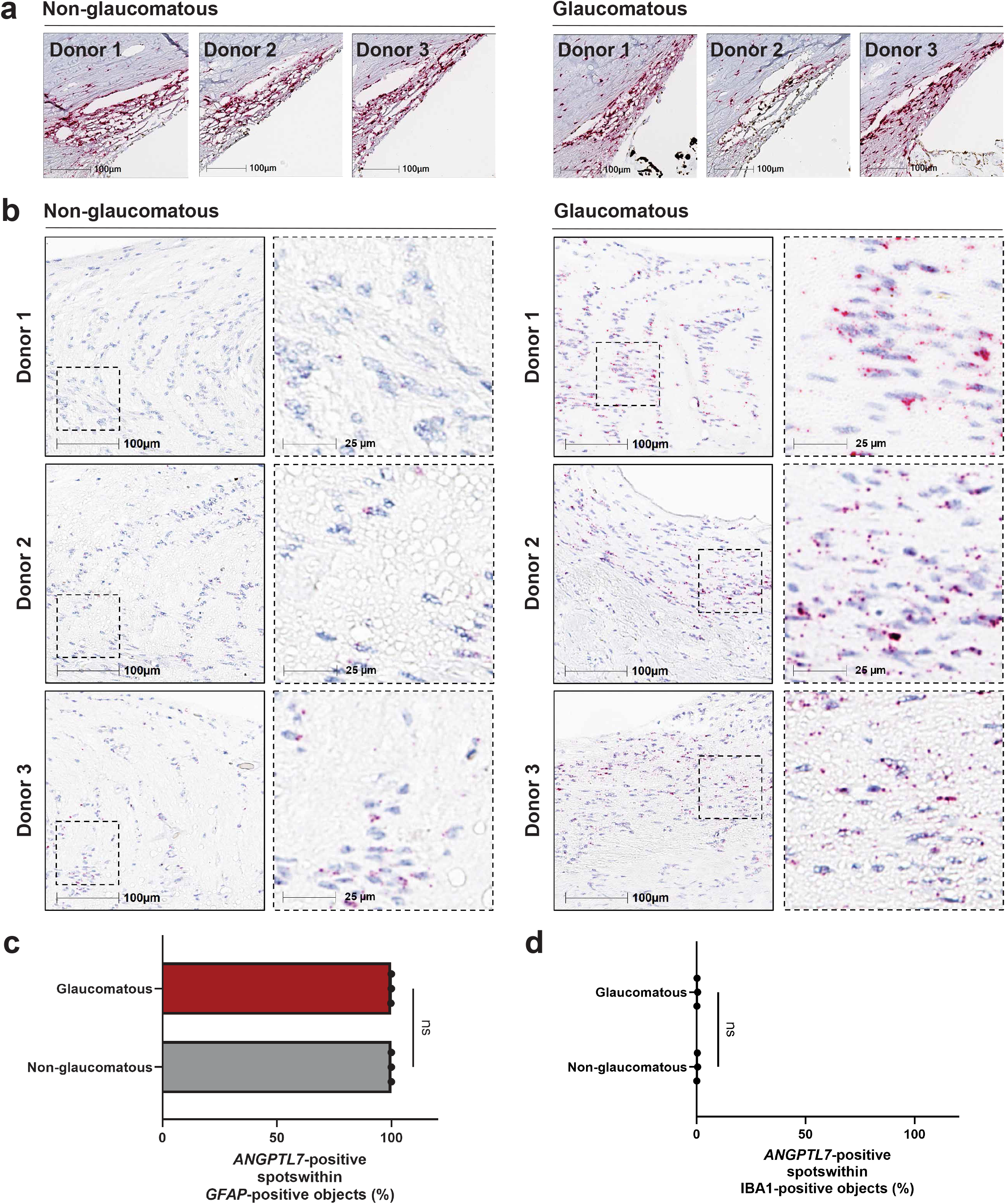
*ANGPTL7* transcript is detected in glaucomatous TM and prelaminar ONHA. Detailed donor information for outflow and ONH chromogenic staining can be found in Supplemental Table 2. **a** Representative images of *ANGPTL7* staining in anterior chamber of non-glaucomatous and glaucomatous human donor tissue by RNAscope. *ANGPTL7* transcript is present in non-glaucomatous and glaucomatous TM region; purple staining indicates *ANGPTL7* mRNA copies. Images selected from n = 4 non-glaucomatous and n = 5 glaucomatous donors. Images from complete donor set are provided in Supplemental Fig. 3a. **b** Representative images of *ANGPTL7* staining in ONH of non-glaucomatous and glaucomatous human donor tissue by RNAscope. Dotted box indicates location of zoomed images (right column for each respective disease state). Images selected from n = 4 non-glaucomatous and n = 5 glaucomatous donors. Images from complete donor set are provided in Supplemental Fig. 3b. **c – d** *ANGPTL7* staining is co-localized almost exclusively with *GFAP* and almost never with IBA1. *GFAP*-positive and IBA1-positive nuclei are classified as *ANGPTL7*-positive or negative and are plotted as a percentage of GFAP or IBA1-associated cells; unpaired t-test for statistical comparison. Detailed donor information for co-staining can be found in Supplemental Table 3.

To confirm expression of ANGPTL7 within astrocytes histologically, we co-stained human tissue sections from non-glaucomatous (n = 3) and glaucomatous (n = 3) post-mortem donors with RNAscope probes against *ANGPTL7* and markers against either astrocytes (*GFAP*; glial acidic fibrillary protein) or mononuclear phagocytes (IBA1; ionized calcium binding adaptor molecule 1). Consistent with chromogenic RNAscope staining, we observe localization of *ANGPTL7* transcript in the prelaminar region of the ONH of normal and glaucomatous tissue sections. *ANGPTL7*-positive spots were almost exclusively associated with *GFAP*-positive cells for normal (99.90 ± 0.13%) and glaucomatous (99.97 ± 0.02%) tissues (Fig. 2c) and rarely associated with IBA1-positive cells for normal (0.20 ± 0.20%) or glaucomatous tissues (0.33 ± 0.11%) (Fig. 2d); data are expressed as average ± standard deviation. These data demonstrate that *ANGPTL7* transcript in the ONH is associated with *GFAP*-positive cells independent of disease status and spatially validates ANGPTL7 colocalization with astrocytes. Our human profiling data finds ANGPTL7 in glaucomatous ONHA, suggesting that the role of ANGPTL7 in glaucoma may be exerted in the back of the eye.

### *Angptl7^-/-^* mice are protected from retinal ganglion cell loss and display reduced glial activation in a glaucoma neurodegeneration model

To evaluate a functional link between ANGPTL7 and neurodegeneration in the posterior eye, we characterized *Angptl7* knockout mice. Given conflicting literature reports regarding IOP phenotypes of *Angptl7^-/-^* mice [11, 12], we conducted our own comprehensive ocular profiling of knockouts and wild-type littermates. We did not observe a statistically significant difference in IOP measurements between 3-month-old *Angptl7^-/-^* mice (11.89 ± 1.01 mmHg) and wild-type littermate controls (11.69 ± 0.77 mmHg) (Fig. 3a). In a longitudinal assessment, IOP was monitored monthly in a separate cohort from 3 to 7 months of age, and no differences were observed between genotypes at any time point (Supplemental Fig. 5). We measured central corneal thickness (CCT) via ocular coherence tomography (OCT) in 3 and 5 month old mice to assess potential correlations between IOP and CCT [28, 29], but observed no significant difference in CCT between genotypes (Fig. 3b, representative images in Supplemental Fig. 6). CCT and IOP results suggest no differences in anterior chamber phenotypes between *Angptl7^-/-^*and wild-type littermate mice.

**Fig. 3.**
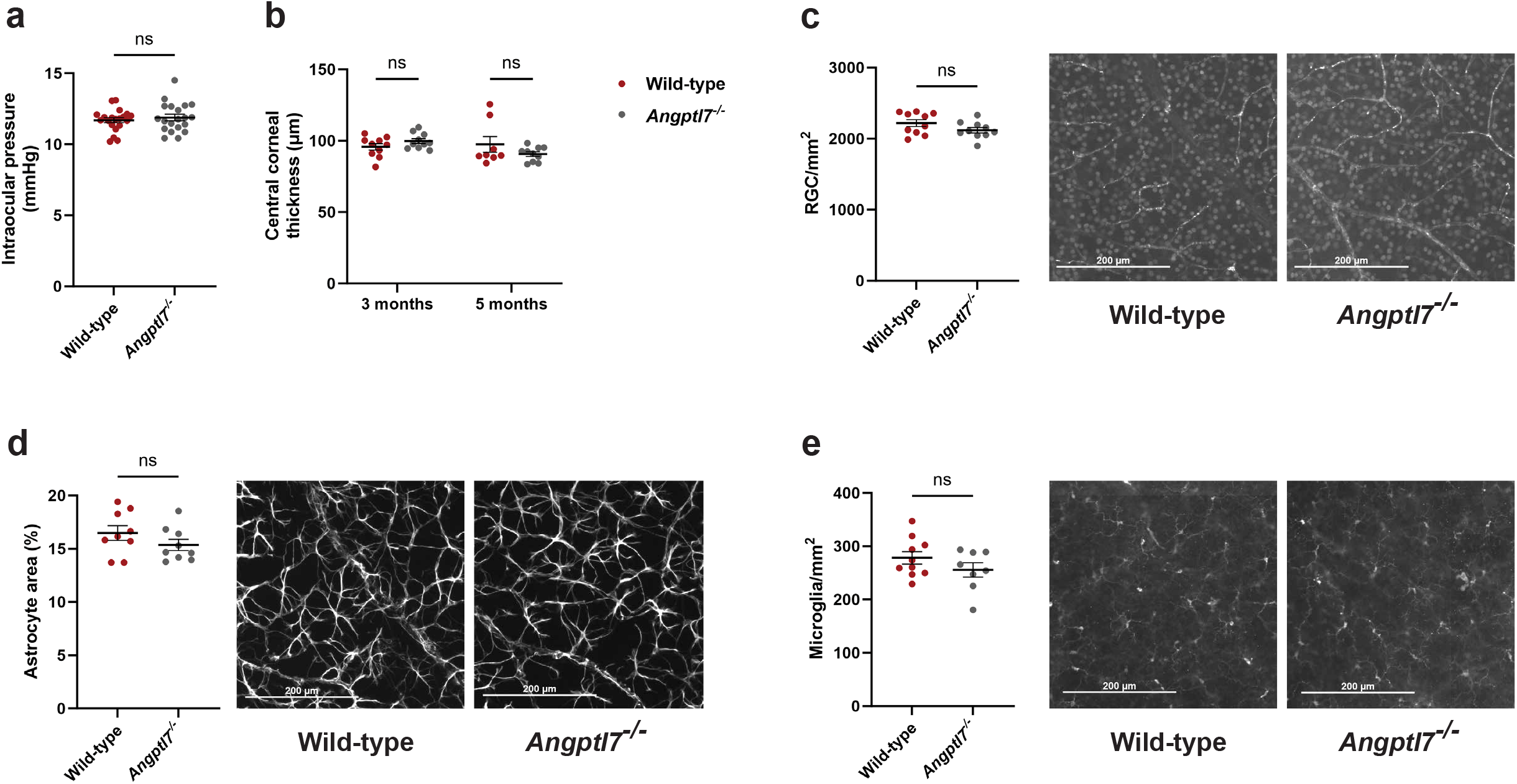
Absence of ANGPTL7 in knockout mice had no impact on ocular phenotypes at baseline. All measurements are expressed as mean ± SEM for respective readouts. Unpaired t-test used for statistical comparison. **a** No significant difference in IOP observed between genotypes at 3 months of age; n = 10 animals per genotype, OD and OS, each plot point represents the mean of ten measurements recorded per eye. **b** No significant difference in central corneal thickness (CCT) was observed between genotypes at 3 or 5 months of age; n = 5 animals per genotype, OD and OS eye. Each plot point represents measurements from individual eyes. **c** No significant difference in RGC counts observed between genotypes. RGCs counts (Brn3a antibody staining) are normalized relative to image area and are reported as RGC/mm^2^. n = 10 animals per genotype, from which one eye was used for profiling. **d** No significant difference in astrocyte measurement observed between genotypes. Astrocytes were counted based on GFAP immunostaining and are normalized relative to image area; data are reported as Astrocyte area (%). n = 9 animals per genotype, from which one eye was used for profiling. **e** No significant difference in microglia counts observed between genotypes. Microglia counts are based on IBA1 immunostaining and are normalized relative to image area; data are reported as Microglia/mm^2^. n = 10 animals for wild-type; n = 8 animals for *Angptl7^-/-^* mice; one eye from each used for profiling. Each plot point in figures c – e represents data from an individual eye.

Glaucoma is an optic neuropathy characterized by RGC death and hallmarks of neuroinflammation such as activation of astrocytes and microglia that likely exacerbate neuronal damage. Given that ONHA express ANGPTL7 and that loss-of-function mutations are protective against development of glaucoma, we asked if *Angptl7^-/-^* mice had detectable differences in RGCs, astrocytes or microglia in neural retina at baseline. Retinal flat-mounts were collected from 3 month old *Angptl7^-/-^* and wild-type littermate mice and stained to quantify RGCs (Brn3a), astrocytes (GFAP) and microglia (IBA1). We observed no significant difference in RGC counts per unit area (Fig. 3c: *Angptl7*^-/-^: 2,119.45 ± 117.06 and wild-type littermates: 2,218.92 ± 151.76), astrocyte area (Fig. 3d: *Angptl7*^-/-^: 15.37 ± 0.53 and wild-type littermates: 16.47 ± 0.68) or microglia density (Fig. 3e: *Angptl7^-/-^*: 255.6 ± 13.57 and wild-type littermates: 278.20 ± 11.60) between genotypes. Qualitatively, there were no obvious differences in glial cell morphology (i.e. soma swelling, retracted/extended processes, etc.). We conclude that loss of ANGPTL7 appears to have no impact on RGC density or glial cell activation under naïve conditions, giving us confidence to initiate a neuroprotection study without concern for confounding elements.

Next, we challenged *Angptl7^-/-^*mice and wild-type littermates in an elevated IOP-induced neurodegeneration model (hyaluronic acid glycidyl methacrylate (HAMA)) [19, 23] to understand whether absence of ANGPTL7 protects from IOP-induced ganglion cell damage or activation of neuroinflammation response. In the HAMA condition (Fig. 4a), IOP was significantly elevated following injection beginning at 3 days and remained significantly elevated at every timepoint until takedown on day 28, observations that were consistent for wild-type and *Angptl7^-/-^*, demonstrating that the elevated IOP phenotype of the model worked as expected in the mutant line. Cumulative IOP, calculated by summing IOP measurements over 28-day time-course for each eye was significantly upregulated in both wild-type and *Angptl7^-/-^* mice and there were no significant differences between wild-type and *Angptl7^-/-^* for naïve or HAMA conditions (Fig. 4b).

**Fig. 4.**
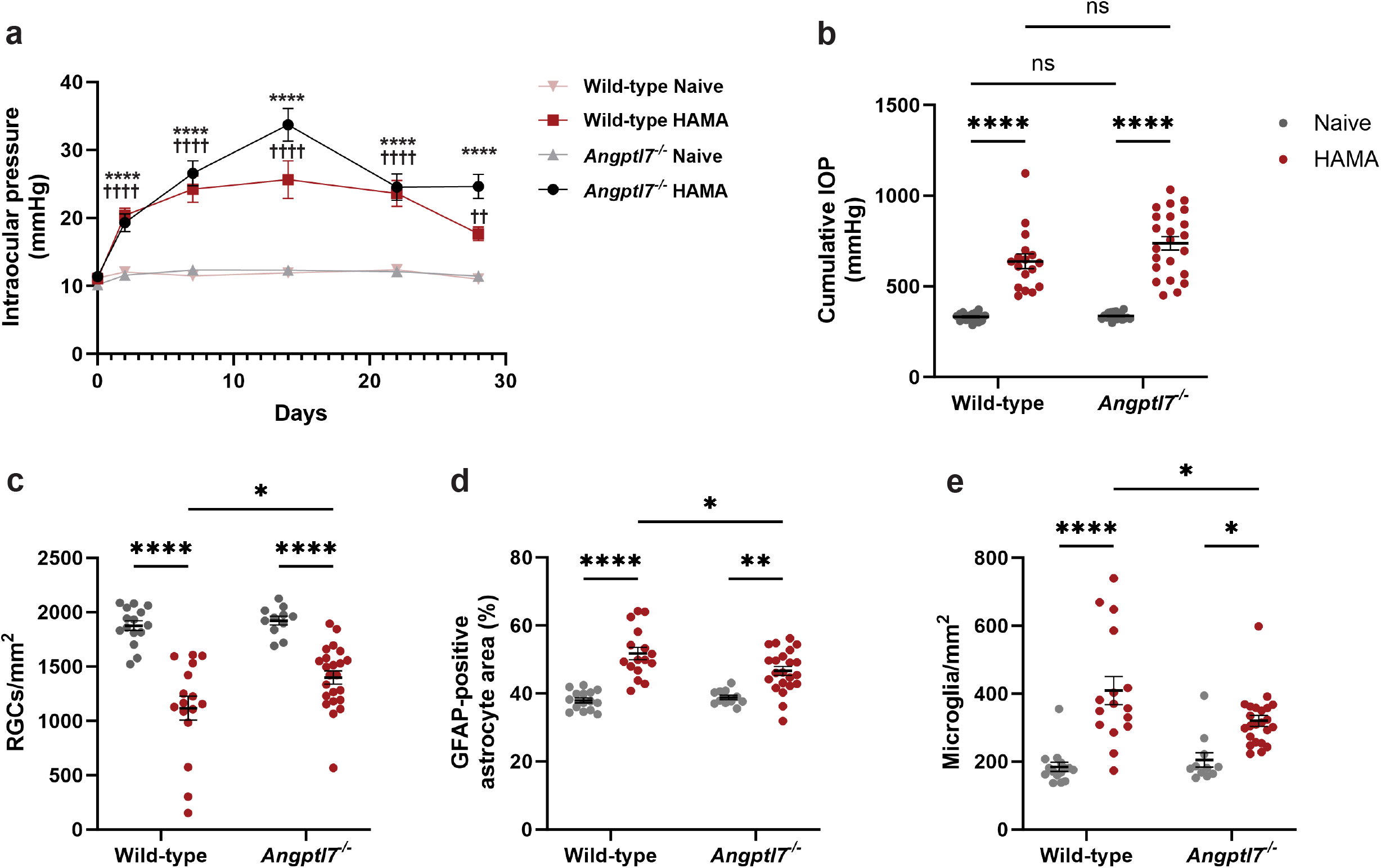
*Angptl7^-/-^*knockout mice exhibit protection from IOP-induced neuronal damage and neuroinflammation in glaucoma neuroprotection model. **a** HAMA injection significantly elevated IOP in *Angptl7^-/-^* and wild-type littermates after 3 days; elevation was sustained at every time-point over the course of 28 days to a level of statistical significance. Each plot point represents an individual eye. One-way ANOVA, Tukey’s multiple comparison test for each time-point. Markers of significance plotted only for comparison of HAMA to wild-type for each genotype; (*) *Angptl7^-/-^* and (^†^) wild-type. Data are represented as mean ± SEM where p > 0.05 is the threshold for significance; ****/^††††^p < 0.0001; ^††^p < 0.01; n = 18 – 23 eyes per group). **b** Cumulative IOP measurement over 28 days shows significant IOP elevation following HAMA injection for both genotypes. Each plot point represents an individual eye. No significant differences were observed between genotypes for naïve or HAMA; n = 18 – 23 eyes per group. c *Angptl7* knockout protects from IOP-induced RGC loss. Data are expressed as RGCs/mm^2^; n = 11 – 23 eyes per group. d *Angptl7* knockout protects from IOP-induced astrocyte activation. Data are expressed as astrocyte area; n = 11 – 22 eyes per group. e *Angptl7* knockout protects from HAMA-induced microglia activation. Data are expressed as microglia/mm^2^; n = 11 – 22 eyes per group. One-way ANOVA, Tukey’s comparison test was used for all statistical analyses in panels b – e; all data are presented as mean ± SEM where p > 0.05 is the threshold for significance; *p < 0.05, **p < 0.01, ****p < 0.0001.

HAMA injection resulted in significant retinal ganglion cell (RGC) loss and increased glial activation in both wild-type (WT) and *Angptl7^-/-^* mice, indicating that the model robustly induced an inflammatory response and glial activation in both genotypes (Fig. 4c-e). However, *Angptl7^-/-^* mice exhibited statistically significant relative protection compared to WT littermates: RGC loss was reduced (27% reduction in *Angptl7^-/-^* versus 40% reduction in WT, providing ∼13% protection; Fig. 4c). Similarly, glial cell activation, as measured by GFAP-positive area, increased in both genotypes following HAMA injection, but *Angptl7^-/-^* mice showed a lower increase (16.8% in *Angptl7^-/-^* vs. 26.6% in WT littermates, representing ∼10% protection; Fig. 4d). Microglia activation was also significantly lower in *Angptl7^-/-^* mice (35.9% increase) compared to WT (54.9% increase), corresponding to 19% protection (Fig. 4e). Collectively, these data demonstrate that while HAMA treatment promotes glial activation and inflammatory signatures in both genotypes, the absence of ANGPTL7 in knockout mice confers statistically significant protection against RGC damage, glial activation, and microglia activation in the glaucoma neuroprotection model, supporting a possible immunomodulatory role for ANGPTL7 within the neural retina.

### Putative ANGPTL7 receptors (LAIR1 and LILRB2) are expressed by ONH immune cells

Members of the ANGPTL family can influence immune cell state through interactions with immunomodulatory receptors containing immunoreceptor tyrosine-based activation (or inhibitory) motifs (ITIMs and ITAMs [30-32]). For example, ANGPTL7 has been shown to interact with the ITIM-domain containing LAIR1 and the ITAM domain-containing LILRB2 when overexpressed [15]. As we observed a reduction in hallmarks of immune cell activation in the *Angptl7^-/-^* HAMA treated group (Fig. 4c-e), we hypothesized a potential link between ANGPTL7 and immune cell signaling. We first confirmed the presence of *LAIR1* and *LILRB2* transcripts within our human snRNAseq dataset. *LAIR1* and (to a lesser extent) *LILRB2* transcripts were enriched in immune cell populations of all tissues profiled from the anterior and posterior eye (Fig. 5a – d). Both genes are expressed in a subset of immune cells (<10% of population) in limbal ring, neural retina and optic nerve, though ONH tissue exhibits a higher percentage of *LAIR1*-positive cells (∼25%). *LAIR1* is enriched in a subset of immune cells in the ONH, the same location where *ANGPTL7* transcript is enriched in astrocytes. RNAscope confirmed the presence of *LAIR1* transcript within the prelaminar ONH region of glaucomatous and non-diseased samples (Fig. 5e). Importantly, these data indicate spatial proximity between ANGPTL7 and LAIR1-positive cells in a manner consistent with a potential interaction.

**Fig. 5.**
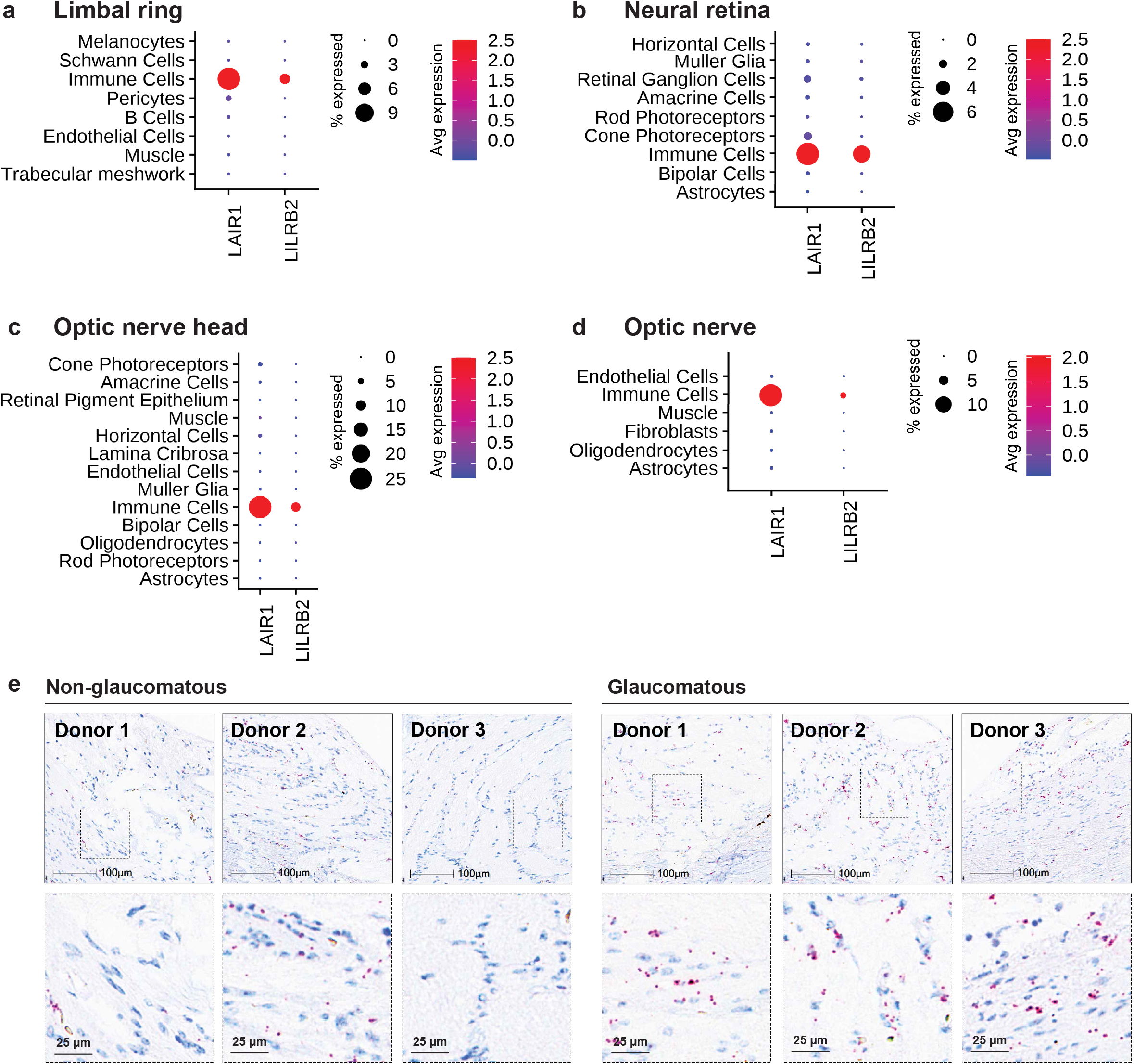
*LAIR1*, *LILRB2* expression in human ocular tissues. snRNAseq was carried out using non-diseased post-mortem human donor material from various ocular tissues; limbal ring (n = 5), macular neural retina (n = 2), ONH (n = 3), optic nerve (n = 3). **a – d** Dot plots showing average expression of *LAIR1* and *LILRB2* transcripts in each cell type. Size of dot indicates percentage of cells expressing transcript and color indicates average number of transcripts. Detailed donor info provided in Supplemental Table 1. **e** Representative images of *LAIR1* staining in ONH of non-glaucomatous and glaucomatous human donor tissue by RNAscope from n = 3 donors for each group. Dotted boxes indicate location of zoomed images (lower panel). Images selected from n = 4 non-glaucomatous and n = 7 glaucomatous donors. Images from complete donor set are provided in Supplemental Fig. 7. Detailed donor information provided in Supplemental Table 4.

### ANGPTL7 binds to LAIR1-expressing cells

To confirm prior reports of ANGPTL7 interactions with LAIR and LILR family members, we transfected U-2 OS cells with cDNAs encoding *LAIR1*, *LILRB2*, and other members of the *LILR* family and added N-Fc ANGPTL7 to the culture. We found that N-Fc ANGPTL7 binds to cells expressing LAIR1 but not cells transfected with *LILR* family members or an unrelated negative control, TEK (Fig. 6a). We then assessed whether ANGPTL7 binds to LAIR1 and LILRB endogenously expressing cells, using THP-1, an immortalized human peripheral blood monocytic line (*LAIR1* and *LILRB2* expression confirmed in Supplemental Fig. 8). Normally cultured in suspension, THP-1 cells can be differentiated into an adherent macrophage-like state when treated with phorbol esters such as PMA [33, 34]. Using this approach, we observed binding of N-Fc ANGPTL7 to the surface of PMA-differentiated THP-1 cells via immunofluorescence (Fig. 6b). ANGPTL7 C-Fc did not bind to PMA-treated THP-1 cells, suggesting that binding of ANGPTL7 likely occurs via the C-terminus, the site of glaucoma protective mutations [6, 11]. N-Fc ANGPTL7 also bound to naïve THP-1 as demonstrated by FACS analysis (Fig. 6c). No binding of mouse Fc alone or N-Fc ANG2 to LAIR1 was observed. Together, these data further establish that ANGPTL7 interacts with LAIR1 and raises the possibility that such interactions mediate glaucoma progression in part through immunomodulatory signaling.

**Fig. 6.**
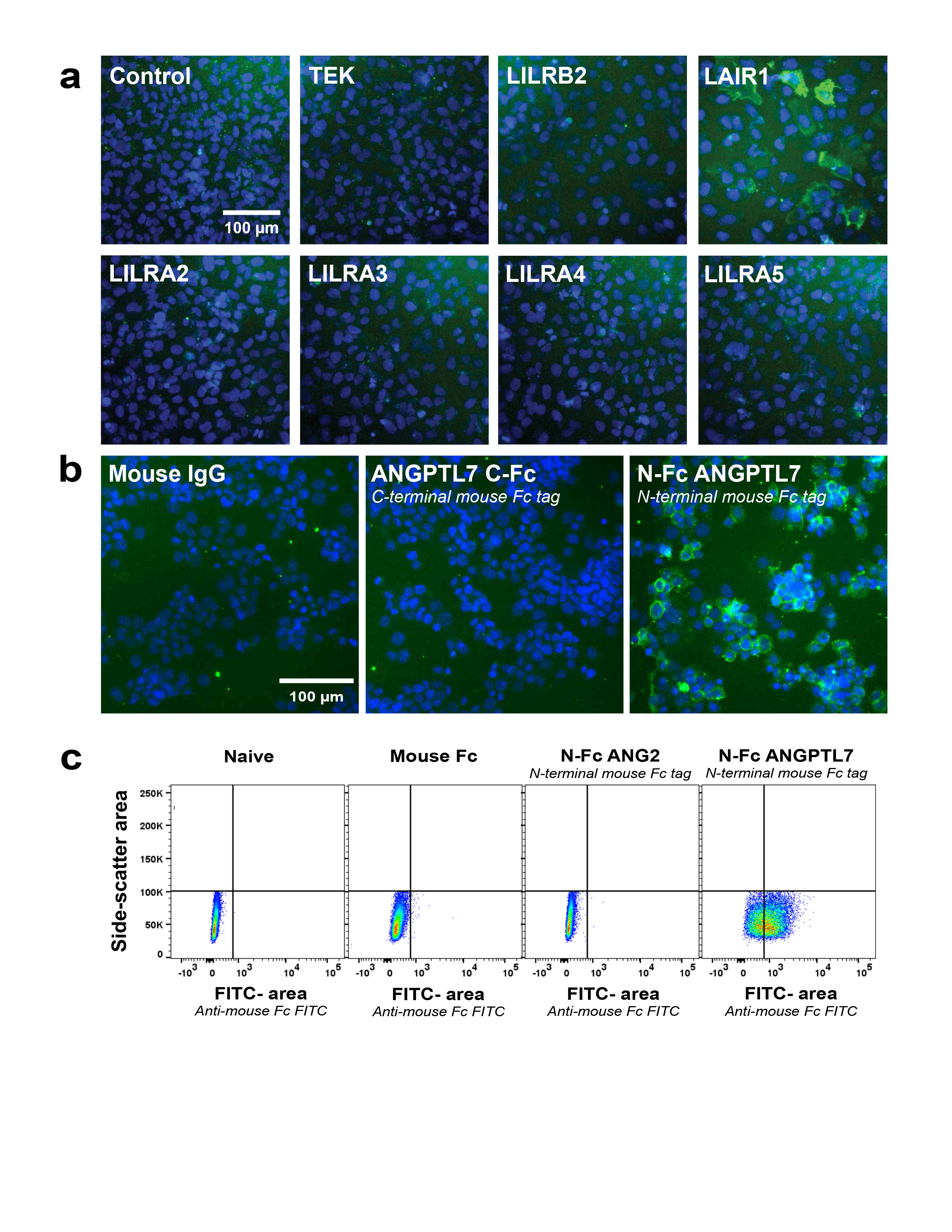
ANGPTL7 binds to LAIR1 positive cells. **a** N-Fc ANGPTL7 binds to LAIR1 over-expressing U-2 OS cells, but not to cells over-expressing other LILR family proteins or negative control (TEK). Green staining indicates detection of mouse Fc tag with anti-mouse Fc (FITC-conjugate); Hoechst dye (blue) indicates nuclear stain. **b** PMA-differentiated THP-1 cells incubated with N-Fc ANGPTL7 or ANGPTL7 C-Fc for 1 hour, then stained with anti-mouse Fc (FITC-conjugate). Representative field of view from n = 4 replicate wells per condition. Green staining indicates detection of mouse Fc; Hoechst dye (blue) indicates nuclear stain. **c** N-Fc ANGPTL7 binds to THP-1 cells by FACS, whereas mouse Fc and N-Fc ANG2 do not.

### Oligomerized ANGPTL7 promotes proinflammatory signaling and attachment in THP-1 cells

ANGPTL7 has been reported to interact with LAIR1 *in vitro* [15], but the downstream consequences of these interactions are unclear. Collagens and complement component 1q (C1Q) signal through LAIR1 to down-regulate tumor necrosis factor alpha (TNF) and interleukin 6 (IL6) [35], but LAIR1 signaling with ANGPTL7 as a ligand had not been evaluated. We hypothesized that ANGPTL7 binding to LAIR1 could stimulate a pro-inflammatory milieu, so we assessed changes to TNF and other cytokines following administration of ANGPTL7 to THP-1 cells. Oligomerization of angiopoietin (ANGPT1/2) and angiopoietin-like proteins (ANGPTL) is often required for signaling activity [36-39], so we used His-tagged rhANGPTL7 crosslinked with an anti-His antibody to create oligomerized protein for functional studies. ANGPTL2, a pro-inflammatory ANGPTL family member, was used as a positive control (without crosslinking), which upregulated *TNF* transcript, as expected [40] (Fig. 7a, middle panel). THP-1 cells were treated with a dilution series of His-tagged rhANGPTL7 (300, 900 & 2,700 ng/mL) with or without anti-His antibody-mediated oligomerization for 48 hours and analyzed for transcriptional changes via qPCR. Oligomerized rhANGPTL7 up-regulated transcription of *IL1RN*, *TNF* and *IL1B* in a concentration dependent manner, which was not observed in cells treated with either His alone or non-oligomerized ANGPTL7 (Fig. 7a).

**Fig. 7.**
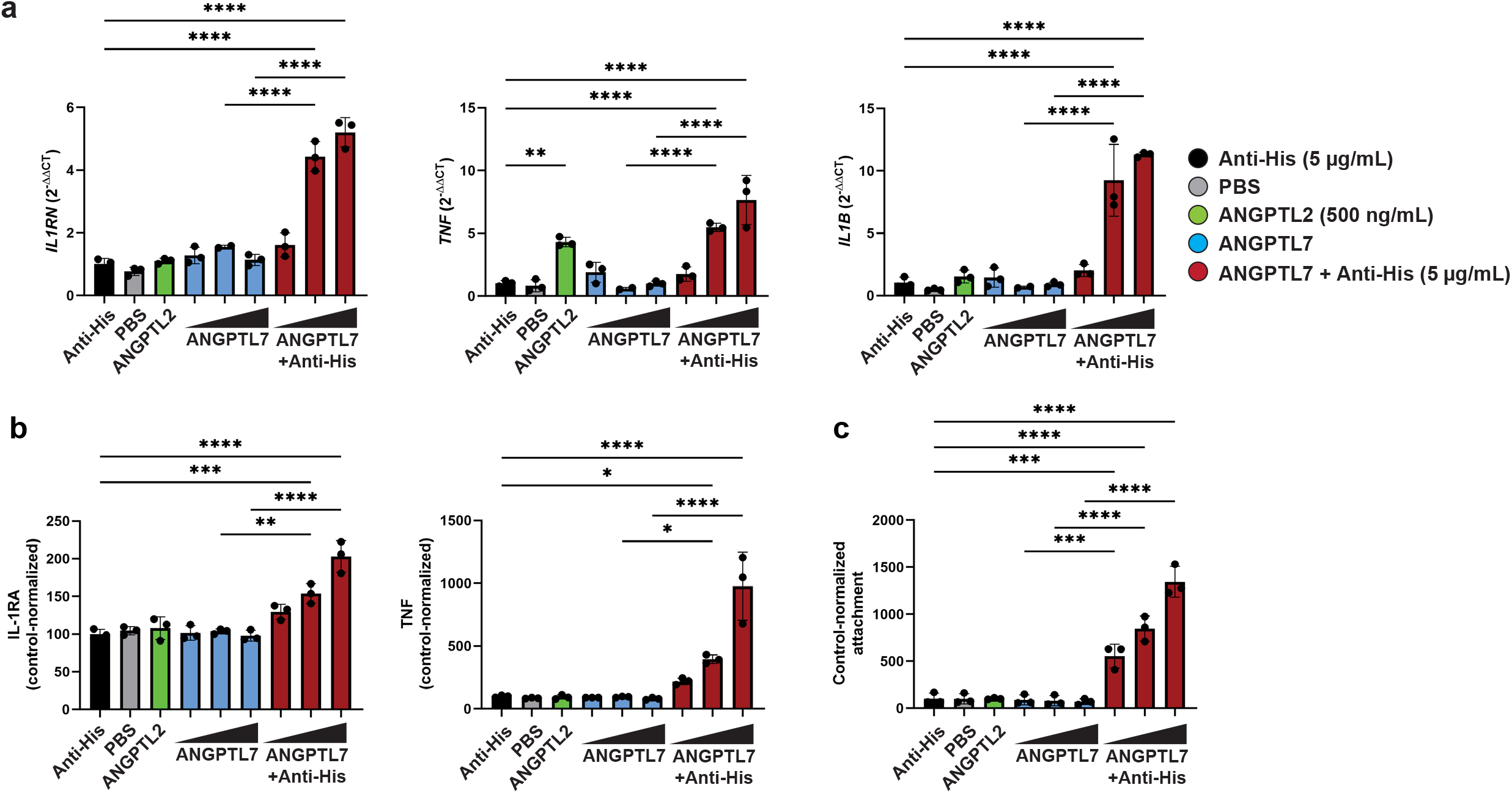
Oligomerized ANGPTL7 activates proinflammatory signaling and attachment in THP-1 cells. Plots are representative of n = 3 experimental replicates, for which figures from other replicates are provided in Supplemental Fig. 11 – 13. Individual plot points represent technical replicates (n = 3) within each experiment. One-way ANOVA, Tukey’s comparison test was used; data are presented as mean ± SEM where p > 0.05 is the threshold for significance; *p < 0.05, **p < 0.01, ***p < 0.001, ****p < 0.0001. **a** Oligomerized ANGPTL7 promotes transcription of *IL1RN*, *TNF* and *IL1B* in a concentration-dependent manner (48 hours). No changes are observed with non-oligomerized ANGPTL7. Data are normalized relative to housekeeping gene (*GAPDH*) and again relative to control treatment (Anti-His) and are expressed as 2^-ΔΔCT^. **b** Oligomerized ANGPTL7 promotes secretion of IL-1RA and TNF into conditioned media in a concentration-dependent manner (96 hours). No changes are observed with non-oligomerized ANGPTL7. Cytokine concentration in conditioned media is based on standard curves and are reported as concentration (pg/mL); all values were in range of standard curve. Data are normalized relative to anti-His control treatment, set to 100. **c** Oligomerized ANGPTL7 promotes attachment of THP-1 cells in a concentration-dependent manner (72 hours). No changes are observed with non-oligomerized ANGPTL7. CTG luminescence data are normalized relative to control (Anti-His) which is set to 100.

These findings were further confirmed at the protein level, via a multiplexed Luminex assay measuring secreted cytokines in conditioned media. Consistent with transcript data, oligomerized ANGPTL7 promoted production and secretion of IL-1RA (*IL1RN* protein) and TNF in a concentration-dependent manner after 96 hours, whereas non-oligomerized ANGPTL7 had no impact on cytokine secretion (Fig. 7b); IL1B protein was measured, but was below the limit of detection of the assay. During conditioned media collection for Luminex analysis, we observed attachment of THP-1 cells to cell culture plates following treatment with oligomerized ANGPTL7. To quantify this observation, we measured cellular ATP content as a proxy readout for cell attachment. Oligomerized ANGPTL7 promoted attachment of THP-1 cells in the range of 72 – 96 hours, with peak attachment observed at 72 hours (Fig. 7c, Supplemental Fig. 13b). Attachment was not observed with non-oligomerized ANGPTL7. These data suggest that oligomerized ANGPTL7 signals through immune cells to promote pro-inflammatory signaling and attachment.

Collectively, our data indicate that ANGPTL7 is expressed by glaucomatous ONHA and that absence of this protein in knockout mice protects from RGC damage and glial activation in a glaucoma neurodegeneration model. Consistent with these findings, ANGPTL7 likely signals through immune cells to initiate proinflammatory signaling, requiring oligomerization for activity. These data suggest a role for ANGPTL7 in immunomodulation of the posterior eye in glaucoma.

## Discussion

To date, exploration of the roles of ANGPTL7 in the eye has focused on its potential regulation of IOP due to high expression in the anterior eye and associations with IOP [6, 9-12]. Our findings expand the role of ANGPTL7 in glaucoma to the posterior eye and implicate a potential direct association with neurodegeneration, with three main lines of evidence. First, ANGPTL7 is expressed in astrocytes of the prelaminar ONH region in human donor eye tissues. Second, we show that loss of ANGPTL7 in knockout mice attenuates IOP-induced RGC injury and glial activation in a glaucoma neurodegeneration model. LAIR1, a proposed receptor of ANGPTL7 is expressed by resident immune cells of the ONH and is present in the prelaminar region of glaucomatous ONH. Third, ANGPTL7 elicits concentration-dependent proinflammatory signaling and substrate adhesion in immortalized monocytic cells (THP-1), with oligomerization required for activity. Together, these findings are consistent with a model in which astrocyte-derived ANGPTL7 modulates immune cell activation within the ONH, contributing to RGC degeneration (Fig. 8).

**Fig. 8.**
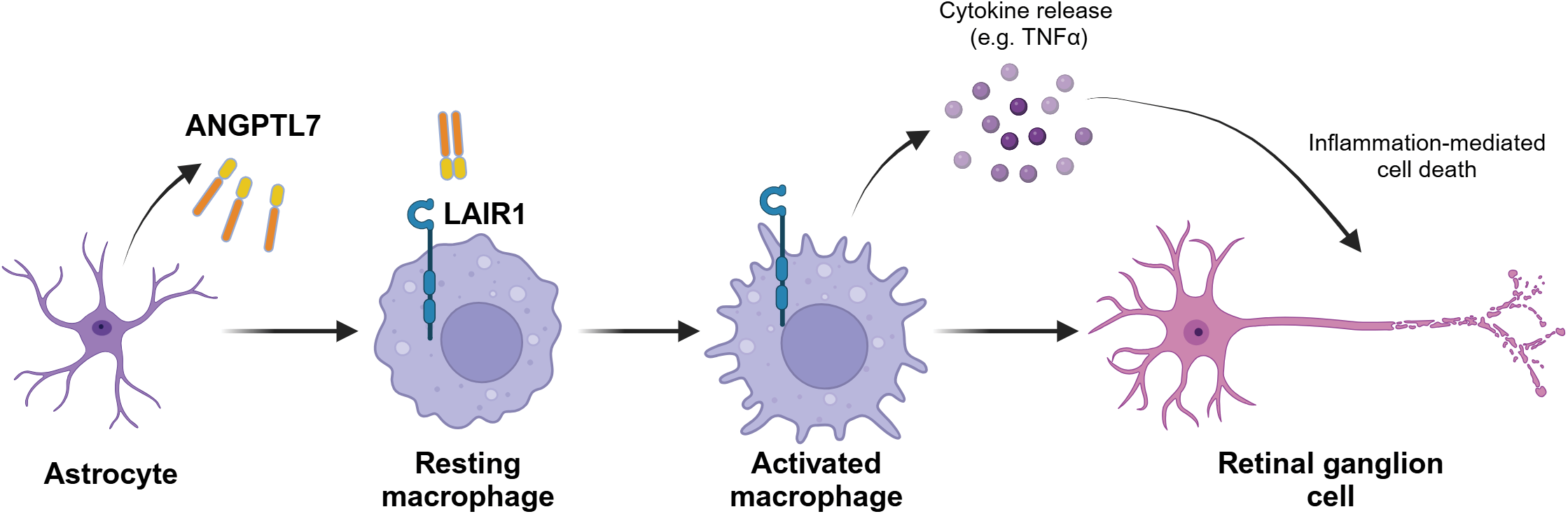
Proposed ANGPTL7 mechanism of action. ANGPTL7 secreted from ONHA activates immune cells, promoting a proinflammatory signaling cascade that activates astrocytes and microglia, eventually leading to retinal ganglion cell damage. Mutant ANGPTL7 variants are not secreted by ONHA and therefore do not activate proinflammatory signaling cascade, preserving retinal ganglion cells, conferring protection against neurodegeneration, progression to glaucoma.

GWAS data linked ANGPTL7 protein-altering variants with lower IOP [6, 11], motivating interest in ANGPTL7 as a potential IOP-modulating target. However, while these variants are associated with protective odds-ratios for glaucoma, the corresponding IOP reductions are modest (<1 mmHg, on average). In some cases, the disease association is substantial, with the Arg220Cys variant conferring an odds-ratio of 0.67 in heterozygotes, and 0.31 in homozygotes [6]. Consistent with this disparity, we observe no differences in IOP between *Angptl7^-/-^* and wild-type mice, nor marked changes in *ANGPTL7* transcript levels in anterior segment tissue from glaucomatous human donors. Instead, we identify robust posterior eye phenotypes across species: *ANGPTL7* transcript is qualitatively elevated in glaucomatous human ONH and in a mouse model of glaucoma-associated neurodegeneration, loss of ANGPTL7 protects against IOP-induced RGC damage and glial/inflammatory cell activation in the neural retina. Together, these findings establish ANGPTL7 relevance in the posterior eye and support a role for ANGPTL7 as a neuroinflammatory contributor to glaucoma. Future studies assessing ANGPTL7 transcript and protein levels across additional glaucoma subtypes, including POAG and normal tension glaucoma, will be important, particularly given that ANGPTL7 is secreted.

Prior studies have reported conflicting effects of *Angptl7* deletion on IOP, with both increased [12] and decreased [11] IOP observed in knockout mice. In contrast, we consistently observed no difference in IOP between *Angptl7^-/-^* and wild-type littermates across multiple cohorts and ages. The *Angptl7^-/-^* mice used in our study were originally generated by Lexicon Pharmaceuticals and subsequently cryo-recovered and expanded by Taconic Biosciences; the same line used by Farber-Katz Brown *et al* [12], who reported increased IOP in knockouts. These mice were generated via homologous recombination targeting exon 1, as described in methods above. By comparison, the *Angptl7* knockout mice used by Praveen *et al* [11], which exhibited decreased IOP, were generated by Regeneron Pharmaceuticals (Tarrytown, NY) using VelociGene and VelociMouse technology, targeting a 571 base-pair region beginning 153 base-pairs upstream of the transcriptional start-site, and maintained on a mixed C57BL/6NTac (63%) and 129SvEvTac (37%) background with embryo transfer into Swiss Webster albino mice. Differences in knockout strategy, genetic background, and colony derivation may therefore contribute to the reported discrepancies in directional IOP changes. Notably, however, published IOP effects across studies are small in magnitude (<2 mmHg, 6 – 11% change), whereas our data robustly demonstrate no genotype-dependent differences in IOP across all examined cohorts. The small magnitude of IOP changes in Angptl7^-/-^ mice are consistent with the modest changes observed in humans with loss-of-function ANGPTL7 variants.

We demonstrate that ANGPTL7 requires oligomerization to initiate signaling; to the best of our knowledge, this is the first report of a functional response driven by oligomerized recombinant ANGPTL7 and establishes a critical consideration for future functional and mechanistic studies. This requirement has important therapeutic implications, particularly for the development of ANGPTL7-targeting antibodies, which must effectively block biologically active multimeric ANGPTL7 rather than merely disrupting oligomer formation. The dependence on oligomerization is consistent with the reported behavior of angiopoietins (ANG) [38, 39, 41] and other ANGPTLs [37, 42], supporting hypotheses [12] that ANGPTL7 functions similarly. ANGPTL family members share a conserved architecture comprising an N-terminal coiled-coil domain implicated in oligomerization and a C-terminal fibrinogen-like domain [43]. We observe receptor binding with N-Fc ANGPTL7, but not with ANGPTL7 C-Fc, suggesting that ANGPTL7 engages its receptor via the C-terminal fibrinogen-like domain, the same region in which reported glaucoma-protective mutations are exclusively localized [6, 11].

We acknowledge limitations of the HAMA glaucoma model used for *in vivo* studies, as this and other microbead-based models represent acute IOP elevation and may not fully capture the chronic nature of human glaucoma. Extending these findings to pressure independent neurodegeneration paradigms (i.e. optic nerve transection or crush) in *Angptl7*^-/-^ mice would further clarify the contribution of ANGPTL7 loss to posterior eye neuroprotection. In addition, our data support the hypothesis that glaucoma-protective ANGPTL7 variants represent loss-of-function alleles and that genetic deletion phenocopies these mutations. Direct testing of this hypothesis, by introducing protective ANGPTL7 variants into the knockout background, would determine whether these mutations confer similar neuroprotection in the HAMA model or exert cell-intrinsic effects on astrocyte function. It is intriguing to speculate that non-secreted or functionally impaired ANGPTL7 variants may shift astrocytes toward a state that is permissive for neuroprotection than neurodegenerative in glaucoma. Such glial/immune cell interactions may operate predominantly in the posterior eye, while potentially overlapping with more modest roles in the anterior segment that influence IOP regulation.

THP-1 cells are widely used for *in vitro* mechanistic studies due to their accessibility, cost-effectiveness and ease of culture; however, as a patient tumor-derived monocytic line, they may incompletely model immune cell behavior *in situ*. More physiologically relevant systems, such as human donor-derived peripheral blood mononuclear cells (PBMC) or inducible pluripotent stem-cell (iPSC)-derived immune cells, may provide additional platforms for elucidating ANGPTL7 signaling and function. The ONH contains resident microglia spanning heterogenous activation states, as well as infiltrating immune populations, and the specific cell types and activation contexts relevant to ANGPTL7 signaling *in vivo* remain unresolved. Our THP-1 studies focused on IL1B and TNF based on prior collagen/C1Q-LAIR1 signaling literature and therefore represent a targeted, hypothesis-driven view of ANGPTL7 activity. Now that the kinetics of oligomerized ANGPTL7-mediated signaling have been established in cultured cells, unbiased transcriptomic approaches, such as bulk RNA-sequencing (bulk RNAseq) in immune cell models could more comprehensively define downstream signaling programs. Notably, complement signaling has been implicated in glaucoma, with elevated C1Q levels reported in subsets of glaucomatous human donor retinas as well as in murine and non-human primate glaucoma models [44], raising the possibility of functional crosstalk between ANGPTL7 and C1Q via LAIR1. In this context, the recent Phase I clinical evaluation of the anti-C1Q monoclonal antibody ANX007 in glaucoma patients [45] provides a unique opportunity to assess whether modulation of complement signaling alters ANGPTL7 levels or activity in ocular tissues or aqueous humor.

ANGPTL7 has been shown to be required for hematopoietic stem cell repopulation [14], supporting a direct role for ANGPTL7 in immune cell regulation and complementing our *in vitro* functional studies and *in vivo* HAMA model observations. Together these findings support a model in which ANGPTL7 acts as an amplifier of neuroinflammation by signaling from astrocytes to resident or infiltrating immune cells within the ONH and neural retina. This signaling promotes the secretion of proinflammatory mediators, initiating a signaling cascade that activates astrocyte and microglia which ultimately leads to RGC degeneration and/or loss. Future studies in *Angptl7^-/-^* mice could focus on longitudinal assessment of transcriptional changes during glaucomatous challenge (e.g., HAMA, optic nerve transection etc.) using bulk sequencing, single-cell RNA sequencing, or spatial transcriptomics to resolve cell-type and region-specific molecular responses.

We propose that under glaucoma-potentiating conditions, ANGPTL7 secreted from ONHA signals through immune cells to drive cytokine upregulation, reinforcing astrocyte and microglial activation and sustaining a neuroinflammatory environment that eventually damages RGCs (Fig. 8). In contrast, glaucoma-protective ANGPTL7 variants, hypothesized to be loss-of-function or effectively ANGPTL7-null, appear to impair ANGPTL7 secretion and thereby prevent immune cell activation, conferring protection against disease progression.

## Author contributions

All authors contributed to reviewing content included in this manuscript. Conceptualization: T.R.V., C.W.W., G.P. Histology and image quantification: C.X., X.W., N.D.K. Mouse experiments were designed/conducted by K.D. and conducted by E.M.C., C.J. Angptl7^-/-^ knockout procurement: M.P. OCT imaging: J.V. In vitro experiments: T.R.V., A.N.G., J.H., I.S.H., J.K., R.G-C., A.C., N.V., A.C. Protein production: C.C.K., L.J.M., A.P.O. Single-nuclei RNA-sequencing analysis: A.B., S-Y.N, C-L.H. RNA-sequencing: L.F., Y.W. Human tissue acquisition/tracking: Y.Q. Writing – original draft: T.R.V. Editing: C.W.W., R.E., V.R., G.P.

## Acknowledgements

We would like to thank Chad Bigelow and Rachida Bouhenni for their support with planning and execution of *in vivo* studies, Rosemarie Cepeda for providing *in vivo* study support, Andrew Krueger for *in vitro* study input. We thank Magali Saint-Geniez & Cynthia L. Grosskreutz for guidance and support. Lastly, we thank the team at Lion’s Eye Institute for Transplant and Research (now Lion’s World Vision Institute) for partnership in procurement of human donor ocular tissue, and the donors and their families for their thoughtful gift to research.

## Supplemental Figures

**Supplemental Figure 1.**
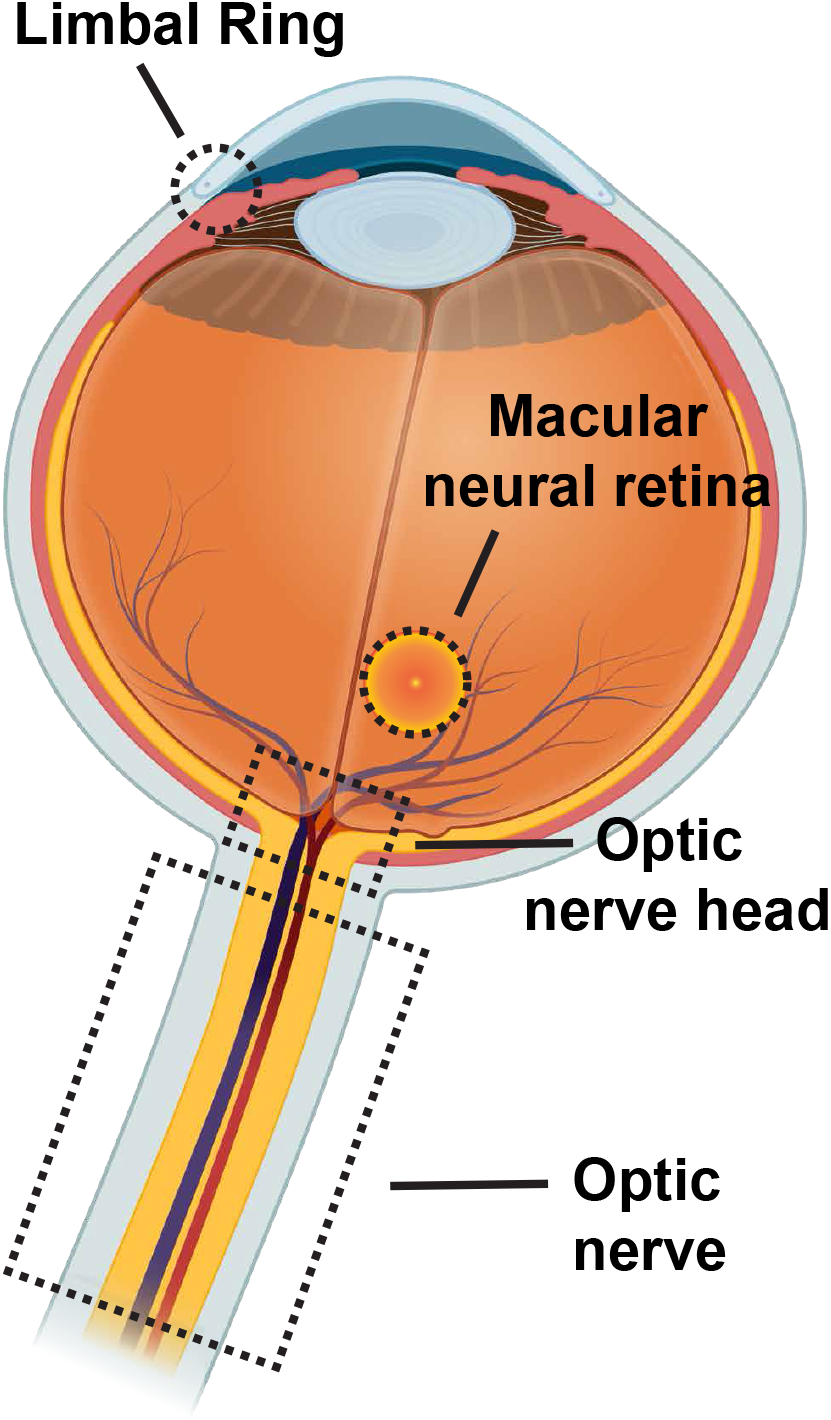
Dissection diagram for snRNAseq of human tissue. snRNAseq was carried out using non-diseased post-mortem human donor material from various ocular tissues. Black dotted regions show locations from where each tissue type was dissected: (1) limbal ring, (2) macular neural retina, (3) ONH, (4) optic nerve.

**Supplemental Figure 2.**
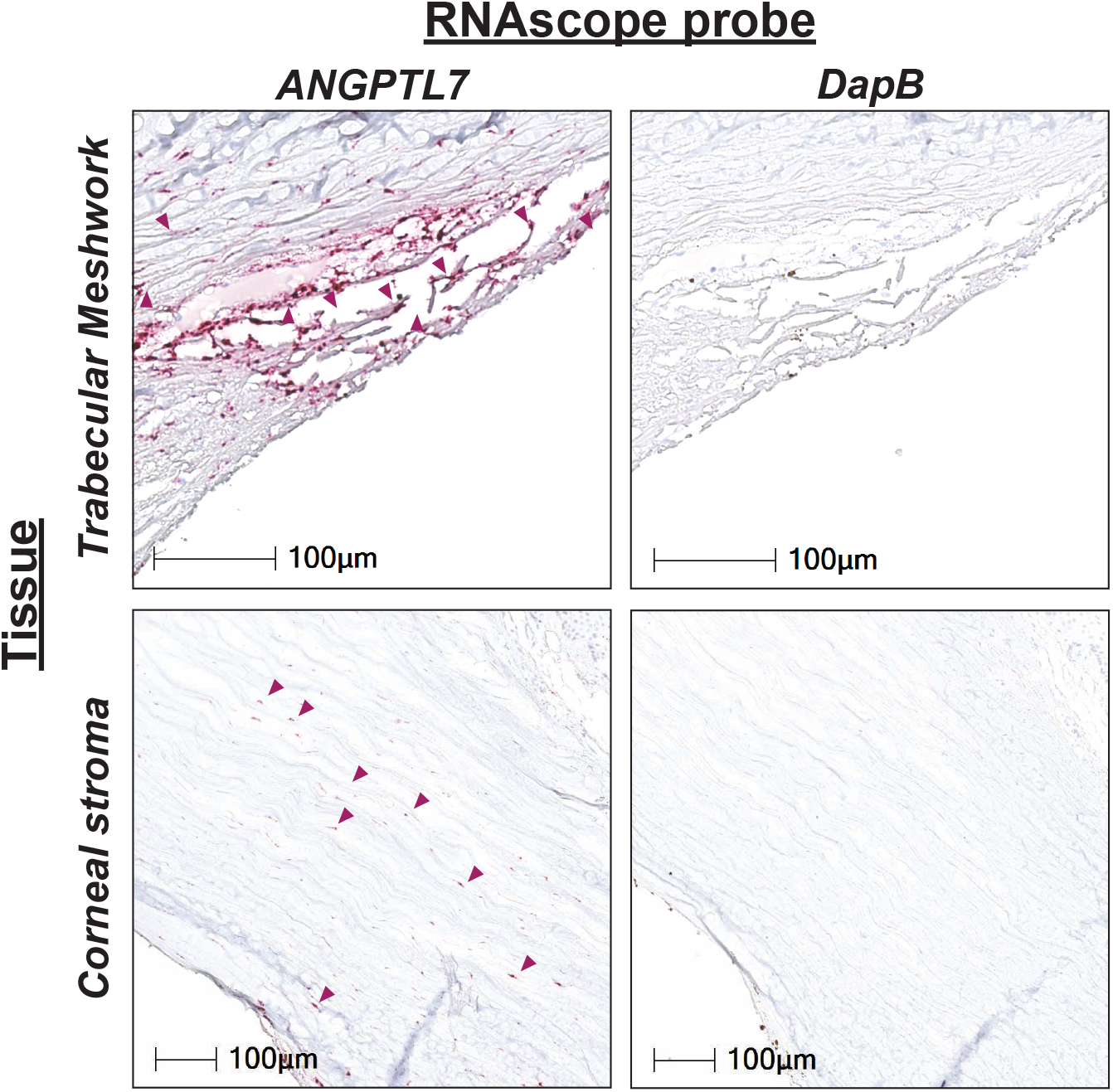
Validation of RNAscope reagents for human tissue staining. Post-mortem human donor tissue stained with *ANGPTL7* and *DapB* (RNAscope). *ANGPTL7* staining present in TM and corneal stroma. Purple arrow indicates transcript detection. Dihydrodipicolinate reductase (*DapB*) is a Bacillus subtilis gene used as a negative control – no staining detected as expected.

**Supplemental Figure 3.**
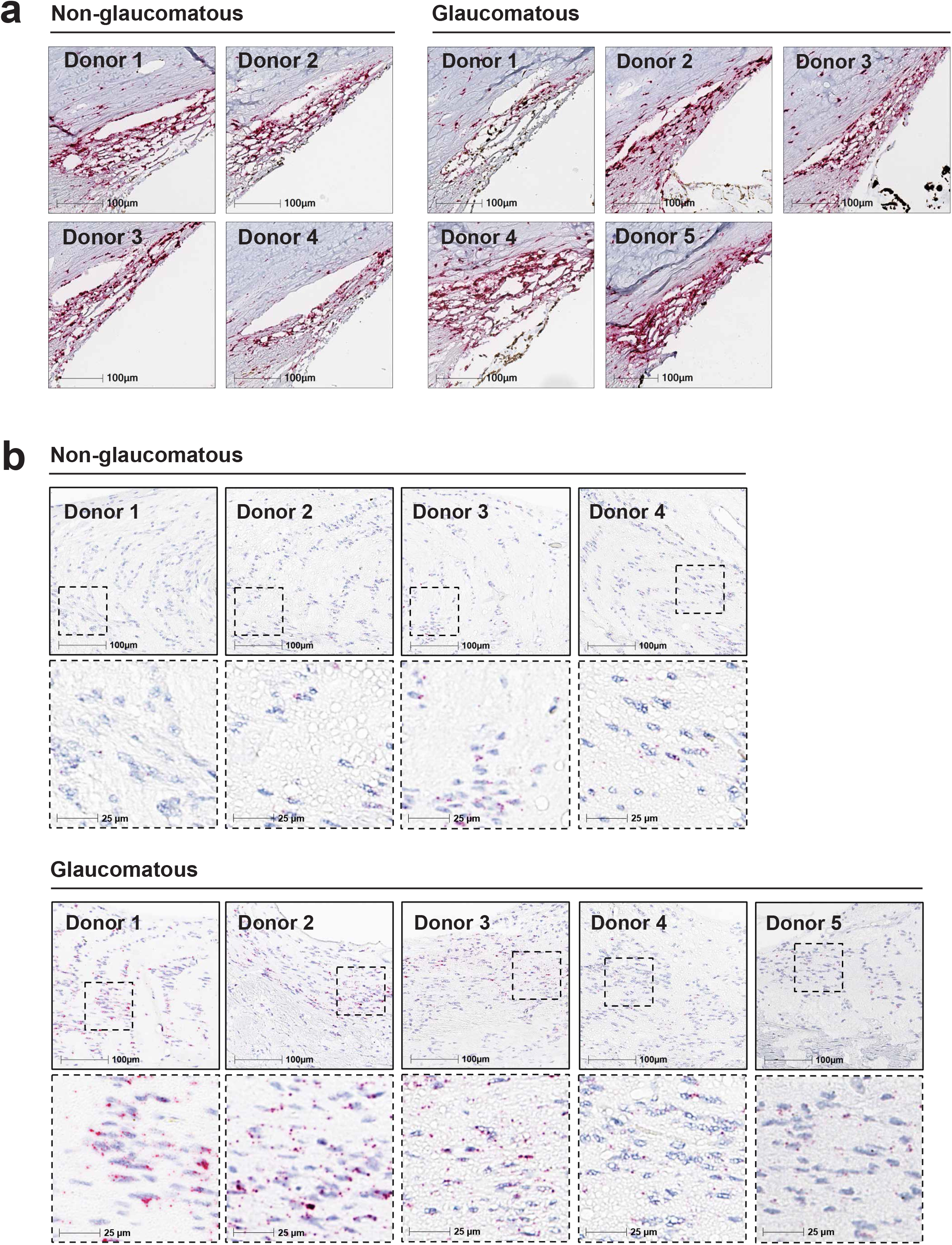
ANGPTL7 transcript in human tissue sections (RNAscope – all samples) **a** *ANGPTL7* RNAscope staining of anterior segment tissues from complete set of donors; non-glaucomatous (n = 4) and glaucomatous (n = 5) donors. **b** *ANGPTL7* RNAscope staining of ONH tissue from complete set of donors; non-glaucomatous (n = 4) and glaucomatous (n = 5) donors. **c** Representative images of *ANGPTL7*-*GFAP*-IBA1 co-staining from a non-glaucomatous (upper panel) and glaucomatous donor (lower panel). Co-staining of *ANGPTL7* (RNAscope), *GFAP* (RNAscope), IBA1 (immunofluorescence) and Hoechst (nuclei stain). **d** Zoomed example of Arivis quantification algorithm. IBA1 or *GFAP*-positive object classification, subsequent counting of ANGPTL7-positive objects within IBA1 (yellow area) or *GFAP* (green area)-positive objects.

**Supplemental Figure 4.**
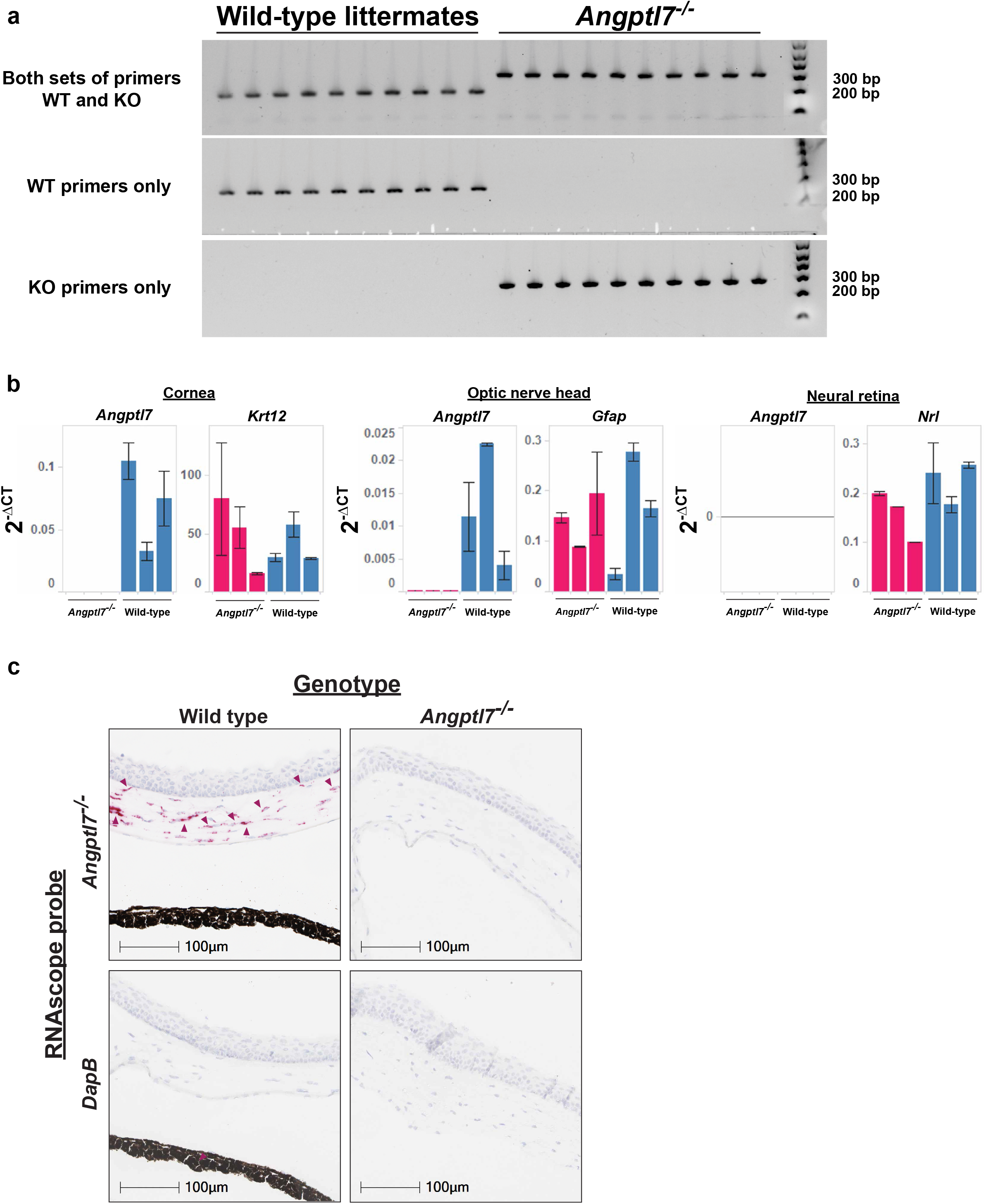
Knockout validation (genotyping, qPCR, RNAscope) **a** Genotyping confirmed knockout status of *Angptl7^-/-^*. Genotyping was performed as described by Lexicon Pharmaceuticals [22]. Tail-snips were collected from 3 month old mice and DNA was extracted using DNeasy Blood & Tissue kit (Qiagen). 10 ng DNA per sample was used as template in PCR reaction. Amplified region of interest was loaded onto a 2% agarose SYBR safe E-gel (Thermo Fisher Scientific) and run for 12 minutes on EG setting. UV images were obtained with a Chemidoc imager (Bio Rad). Expected PCR product sizes for knockouts are 296 base-pairs; expected PCR product sizes for wild-type are 208 base-pairs. Here we confirm reported genotypes. Mutant primer sequences: (1) ACAATATTTAAGATGCTGCC and (2) CCCTAGGAATGCTCGTCAAGA. Wild-type primer sequences: (1) GAGGTACAGCAGCAACAATA and (2) TGTTCCCCTTAGAGTCT. **b** No *Angptl7* transcript detected in cornea, ONH of *Angptl7^-/-^* mice by qPCR. Previously, Regeneron reported enrichment of *ANGPTL7* transcript levels in cornea, sclera, trabecular meshwork and ONH by bulk RNA-seq across multiple species (human, green monkey and mouse); no *ANGPTL7* transcript was detected in neural retina [11]. Eyes from *Angptl7^-/-^* and WT littermates (n = 3 animals per genotype, n = 6 eyes) were enucleated and dissected into (1) cornea, (2) ONH and (3) neural retina (negative control); tissues were homogenized using a Precellys homogenizer after which RNA was extracted using a Direct-zol kit (Zymo Research). An equivalent amount of RNA was used to construct cDNA libraries with SuperScript IV VILO Mastermix (Thermo Fisher Scientific) after which qPCR was performed using TaqMan reagents. Data show 2^-ΔCT^ where delta (Δ) CT equals Experimental CT – housekeeping CT (*Gapdh*). Data are grouped on X axis by animal (1 through 3) for each respective genotype and are presented as median ± median absolute deviation. Additional probes were used as tissue-specific controls; *Gfap* – ONH astrocytes, *Nrl* – neural retina rods, *Krt12* – corneal epithelium. Wild-type cornea and ONHs were *Angptl7*-positive, as expected; *Angptl7* was not detected in these tissues from *Angptl7^-/-^*mice. *Krt12* transcript levels were consistently detected in corneal tissues regardless of genotype; *Gfap* transcript levels were consistently detected in ONH tissues regardless of genotype. *Angptl7* transcript was absent from neural retina, regardless of genotype, as expected. TaqMan probes used in this experiment are detailed in Supplemental Table 7. **c** *Angptl7* transcript absent from *Angptl7^-/-^* cornea (RNAscope) RNAscope was utilized to assess *Angptl7* transcript levels in *Angptl7^-/-^*mice and wild-type littermates. *Angptl7* was present in corneal stroma of wild-type littermates, but absent in knockouts, consistent with genotyping and qPCR data. *DapB* staining used as negative control. Purple arrow indicates transcript detection. Representative images shown from n = 4 animals per genotype.

**Supplemental Figure 5.**
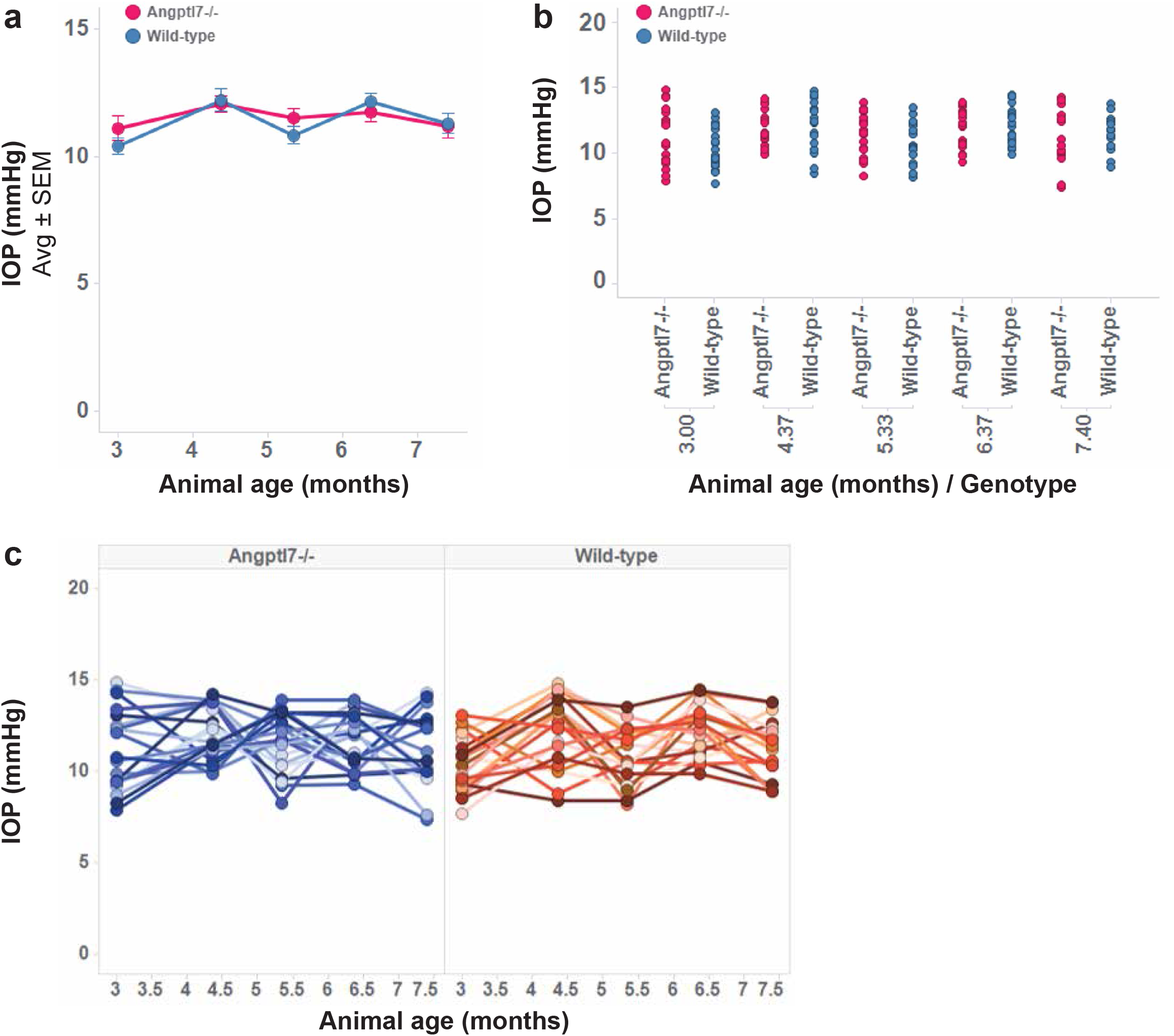
Longitudinal IOP monitoring of *Angptl7^-/-^* and wild-type littermate mice. No significant difference in IOP was observed between *Angptl7^-/-^* and wild-type mice over a 4.5 month timeframe, in animals aged 3 to 7.5 months. There were n = 10 animals per genotype, measurements from OD and OS. One animal from each genotype had to be euthanized before the end of the study, so later timepoints were not recorded for these animals. IOP was monitored every 1 – 1.5 months, measurements are an average of n = 10 repeated measurements. **a** Line plot showing average ± standard error IOP measurements for each genotype. **b** Individual plot points showing no difference in IOP measurement at each individual timepoint. **c** IOP measurement for each individual animal/eye over 4.5 month timeframe.

**Supplemental Figure 6.**
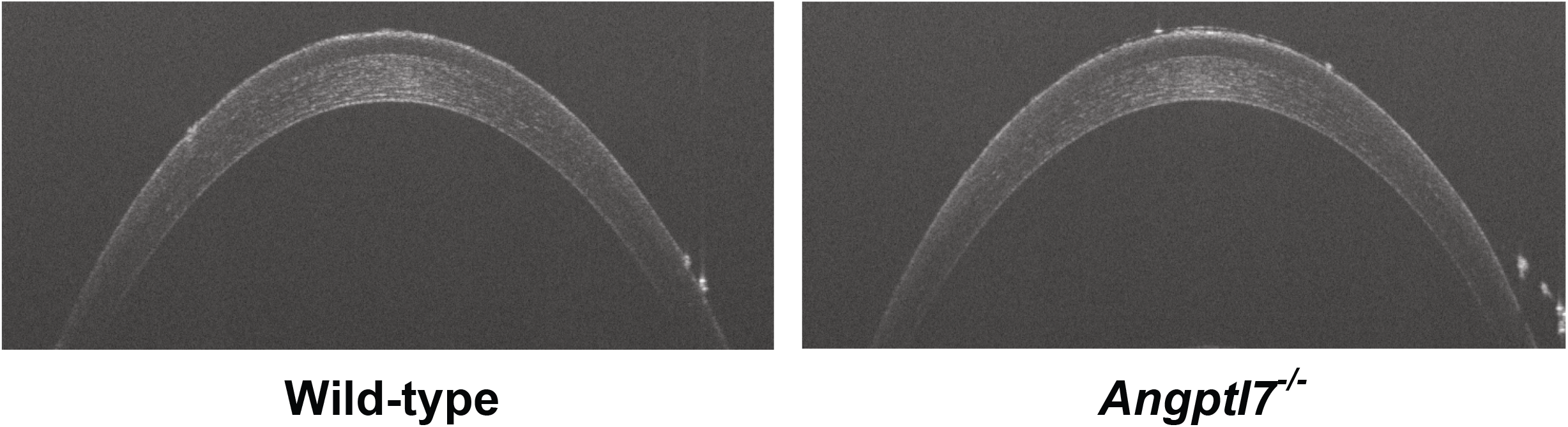
Corneal OCT in *Angptl7^-/-^* and wild-type littermates. Representative images of central cornea OCT from *Angptl7^-/-^* and wild-type littermate mice at 3 months.

**Supplemental Figure 7.**
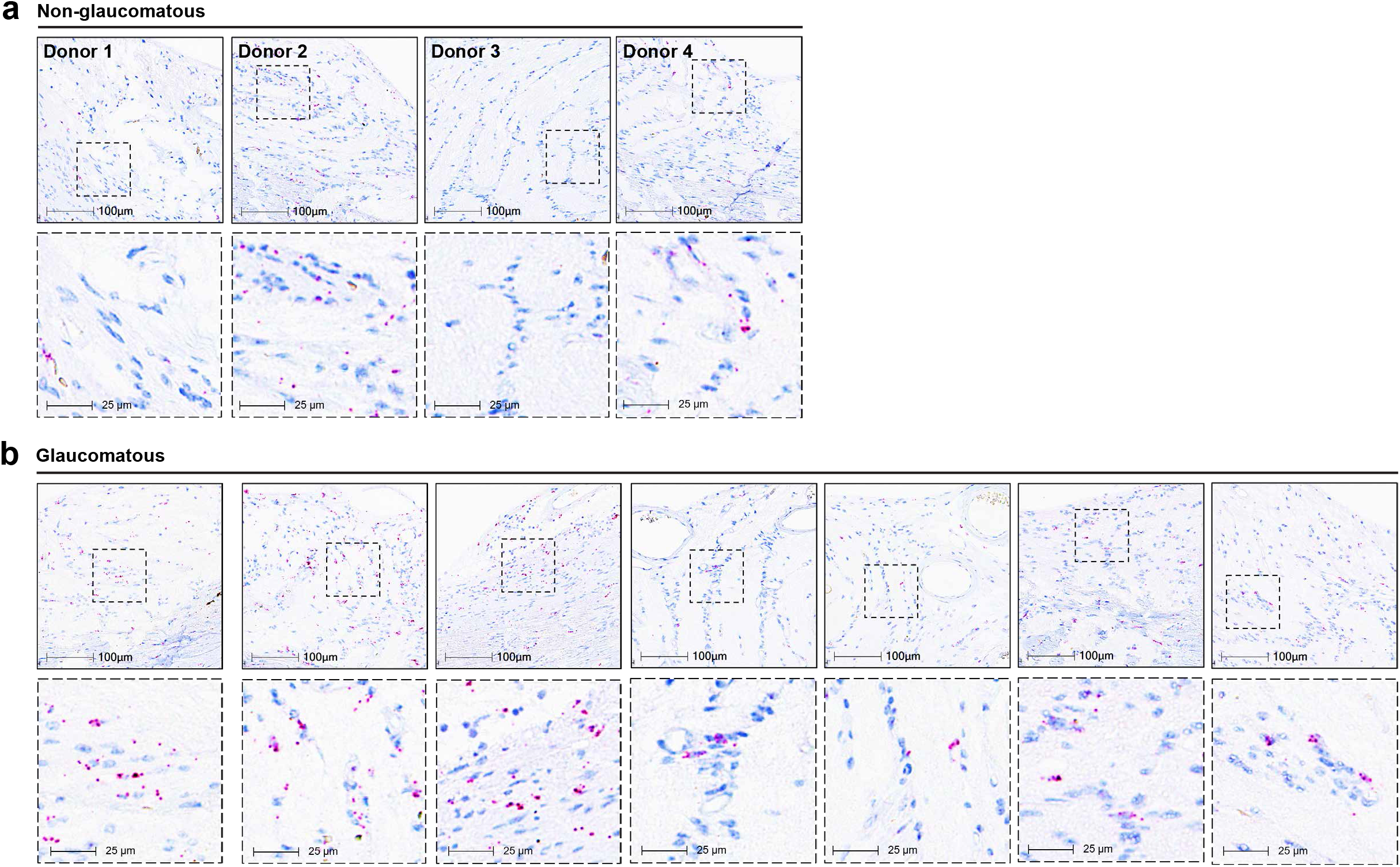
L*A*IR1 transcript in human tissue sections (RNAscope – all samples) *LAIR1* RNAscope staining of ONH tissue from complete set of donors; non-glaucomatous (n = 4) and glaucomatous (n = 7) donors.

**Supplemental Figure 8.**
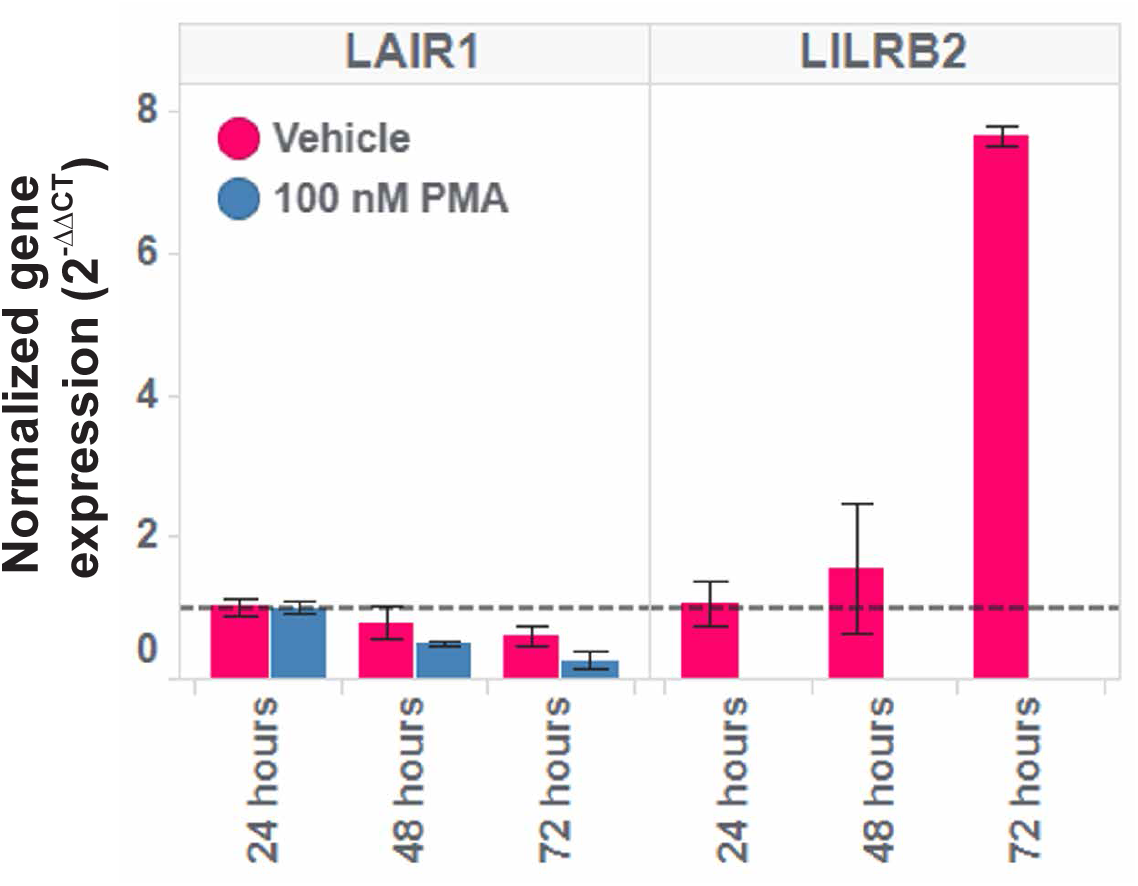
Confirmation of *LAIR1* expression in THP-1 cells (qPCR) *LAIR1* transcript was detected in PMA-treated THP-1 cells by qPCR. Cells were cultured as described in methods. THP-1 cells were treated with 100 nM PMA for 24, 48 and 72 hours, after which RNA was collected, cDNA library was constructed and qPCR was executed using TaqMan reagents against *LAIR1* or *LILRB2* (FAM probe) and *GAPDH* (VIC probe) as a housekeeping gene. Data are expressed as 2^-ΔΔCT^ where ΔCT equals experimental CT (*LAIR1*, *LILRB2*) – housekeeping CT (*GAPDH*) and ΔΔCT equals ΔCT – 24 hour ΔCT. PMA-treated THP-1 cells still express *LAIR1*, but not *LILRB2*. *LAIR1* CT values were below 25 suggesting high level of transcript.

**Supplemental Figure 9.**
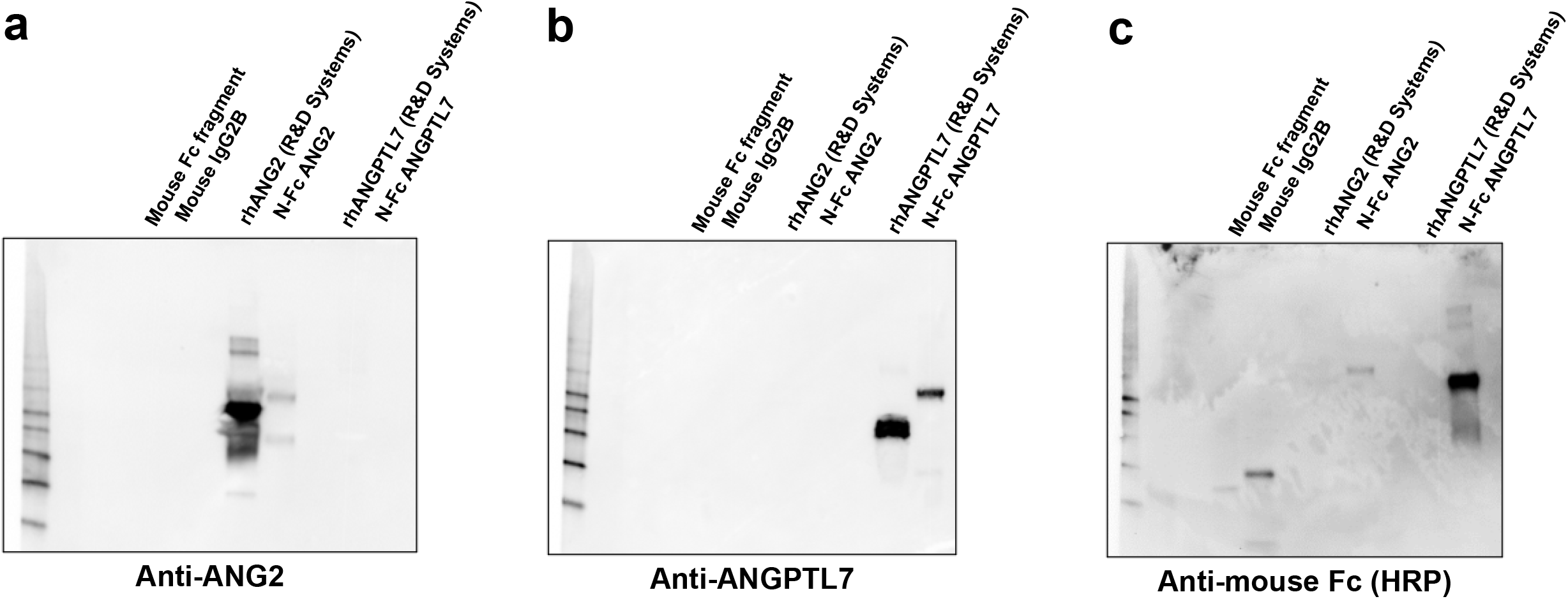
Validation of Fc-tagged proteins (Western blot) **a** Anti-ANG2 antibody (R&D AF623) detects recombinant human ANG2 (rhANG2 – R&D Systems) and Fc-tagged ANG2. Antibody does not detect ANGPTL7 or Fc controls. **b** Anti-ANGPTL7 antibody (R&D AF914) detects recombinant human ANGPTL7 (rhANGPTL7 – R&D Systems) and N-Fc ANGPTL7. Antibody does not detect ANG2 or Fc controls. **c** Anti-mouse Fc detects Mouse Fc fragment and IgG2, as well as Fc fragments on ANG2-Fc and N-Fc ANGPTL7. No signal from untagged rhANG2 or rhANGPTL7 from R&D.

**Supplemental Figure 10.**
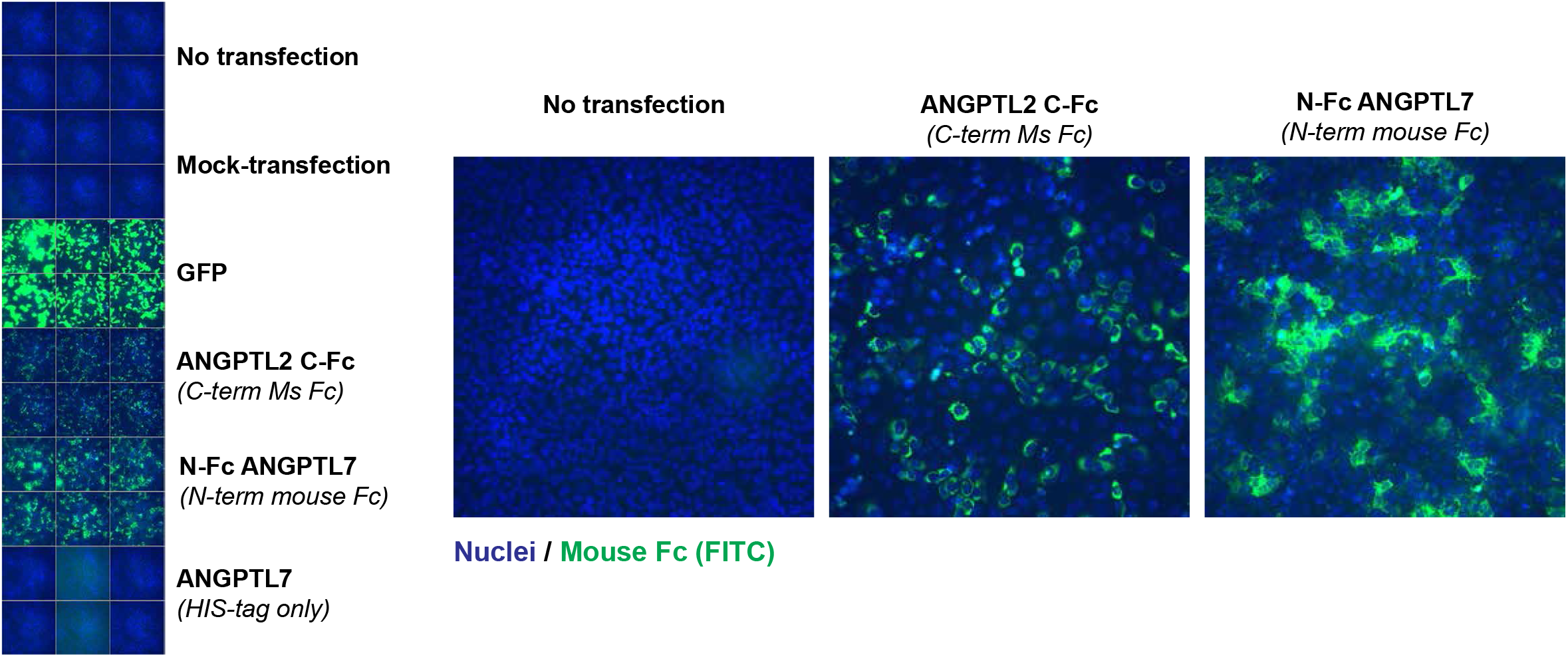
Validation of anti-mouse Fc (FITC) antibody for binding experiments. ANGPTL plasmids containing mouse Ig Fc domain as well as His-tagged ANGPTL7 plasmid were transfected into U-2 OS cells using FuGENE HD reagents (Promega). After 48 hours, cells were fixed with 4% paraformaldehyde, blocked and permeabilized with tris-buffered saline (TBS) supplemented with 10% goat serum and 0.1% Triton X-100 for 1 hour at room temperature. Plates were then stained overnight with anti-mouse Fc (FITC) antibody (Sigma Aldrich, AP127F) at 4 °C according to manufacturer’s instructions. Fluorescent images were collected at 20X magnification using an InCell2200 (Cytivia).

**Supplemental Figure 11.**
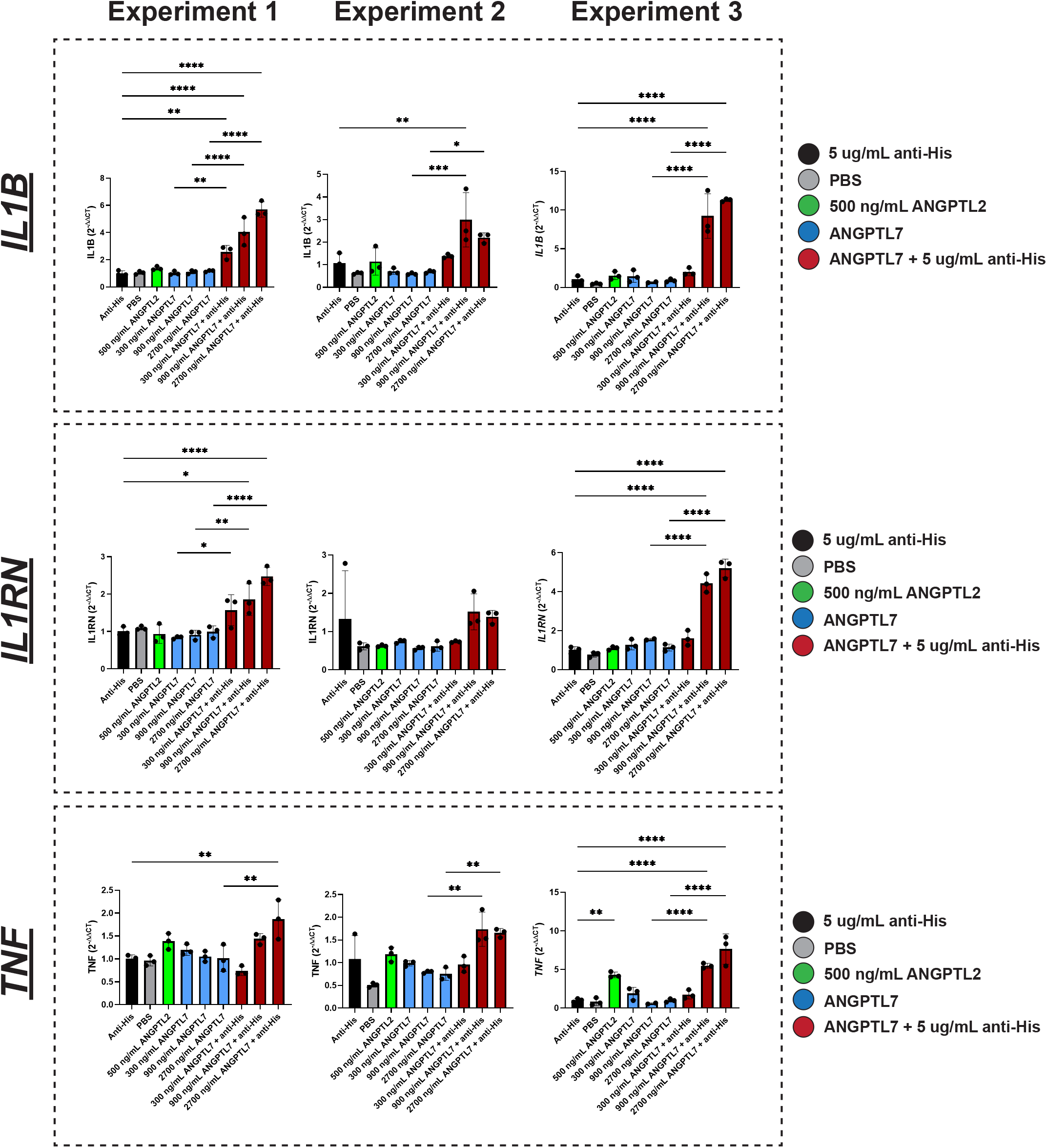
ANGPTL7 upregulates proinflammatory transcripts in THP-1 cells (qPCR experimental replicates) Each column represents experimental replicates (n = 3) and rows are for each transcript profiled. This figure corresponds to Fig. 7a. Oligomerized ANGPTL7 promotes transcription of *IL1RN*, *TNF* and *IL1B* in a concentration-dependent manner (48 hours). No changes are observed with non-oligomerized Angptl7. Data are normalized relative to housekeeping gene (*GAPDH*) and again relative to control treatment (Anti-His) and are expressed as 2^-ΔΔCT^. One-way ANOVA, Tukey’s multiple comparison test was used (mean ± standard deviation; ****p < 0.0001; ***p <0.001, **p < 0.01, *p < 0.05).

**Supplemental Figure 12.**
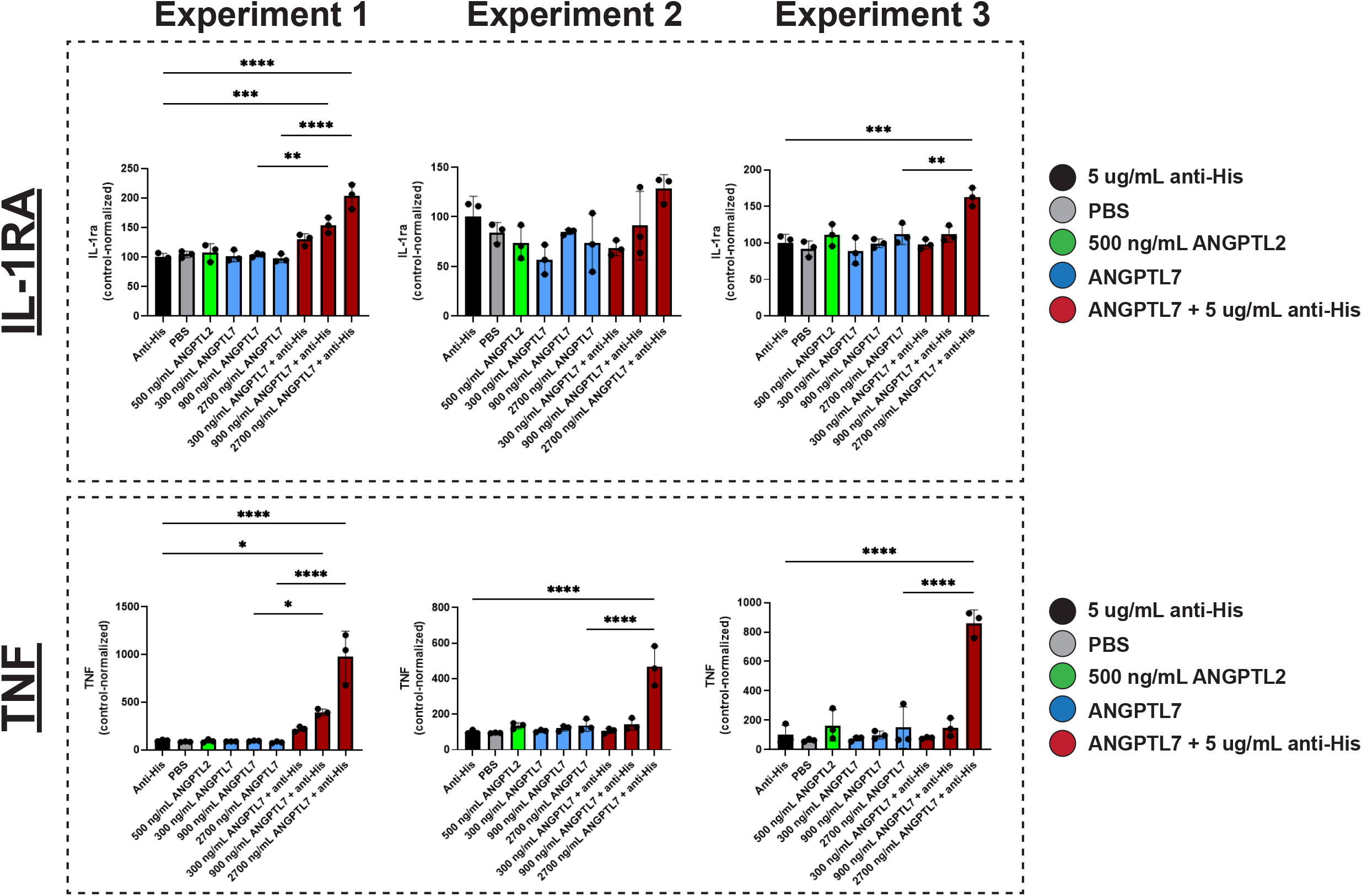
ANGPTL7 upregulates proinflammatory protein secretion in THP-1 cells (Luminex experimental replicates) Each column represents experimental replicates (n = 3) and rows are for each protein measured. This figure corresponds to Fig. 7b. Oligomerized ANGPTL7 promotes secretion of IL-1RA and TNF in a concentration-dependent manner (96 hours). No changes are observed with non-oligomerized ANGPTL7. Cytokine concentration in conditioned media is based on standard curves and are reported as concentration (pg/mL); all values were in range of standard curve. Data are normalized relative to anti-His control treatment, set to 100. One-way ANOVA, Tukey’s multiple comparison test was used (mean ± standard deviation; ****p < 0.0001; ***p <0.001, **p < 0.01, *p < 0.05).

**Supplemental Figure 13.**
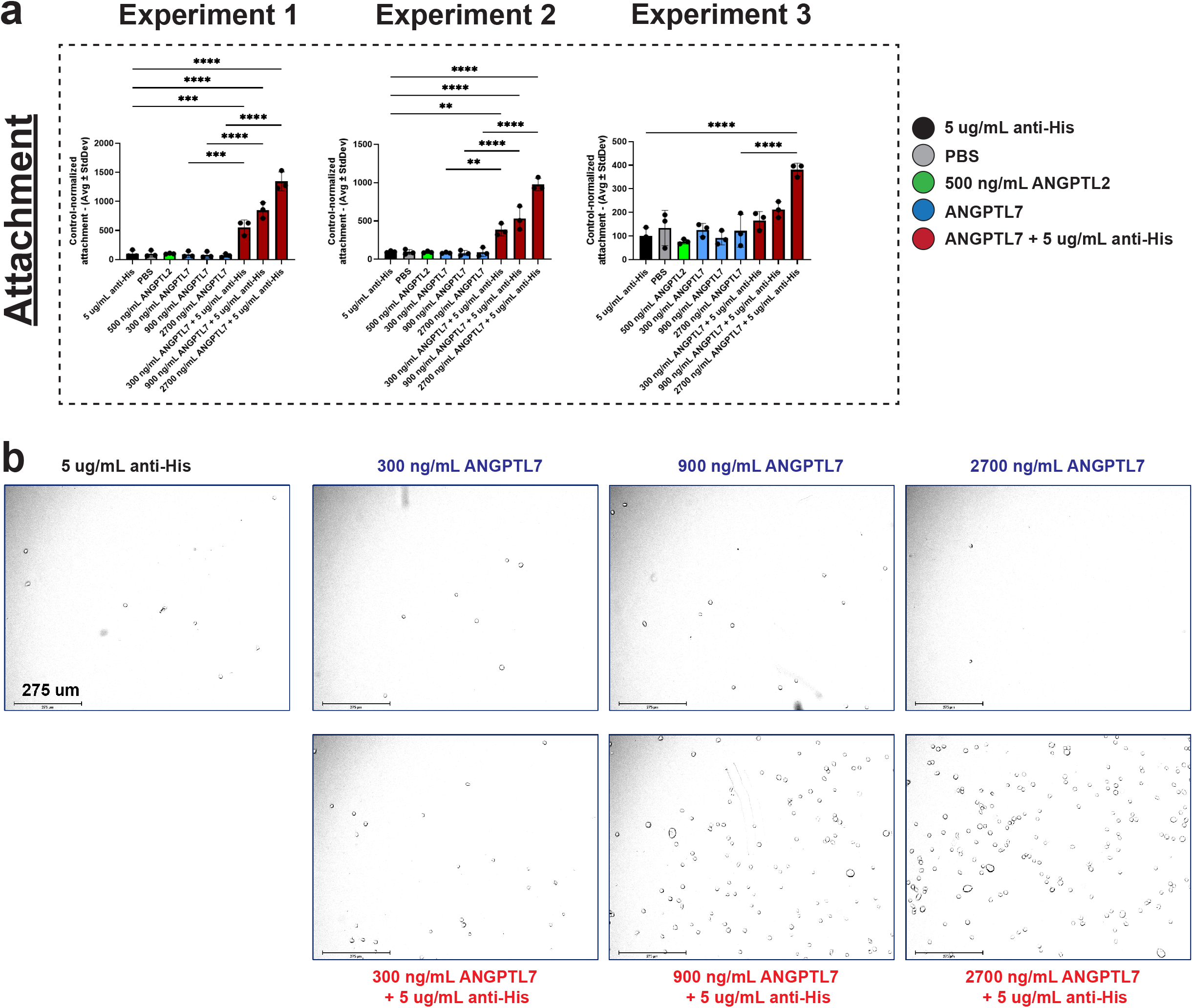
ANGPTL7 upregulates attachment of THP-1 cells (CTG/Attachment assay replicates) **a** Each column represents an independent experimental replicate (n = 3); corresponds to Fig. 7c. Oligomerized ANGPTL7 promotes attachment of THP-1 cells in a concentration-dependent manner (72 hours). No changes are observed with non-oligomerized ANGPTL7. CTG luminescence data are normalized relative to control (Anti-His) which is set to 100. One-way ANOVA, Tukey’s multiple comparison test was used (mean ± standard deviation; ****p < 0.0001; ***p <0.001, **p < 0.01, *p < 0.05). **b** For attachment assay, THP-1 cells were plated in 96-well plates at a density of 70,000 cells per well. Cells were treated for 72 hours. Cell suspension/supernatant was removed for cytokine analysis via Luminex. Adherent plates were washed twice with PBS before addition of CTG 2.0 reagent (Promega, Madison, WI). Representative images taken post-PBS wash, pre-CTG 2.0 addition. Oligomerization of ANGPTL7 is required to promote attachment of THP-1 cells in a concentration-dependent manner.

**Supplemental Table 1.** Human donor information (snRNAseq) – non-glaucomatous donors.

| Tissue | Donor | Age | Sex | Race | AMD | Death-to-preservation time (hours:minutes) |
| --- | --- | --- | --- | --- | --- | --- |
| Limbal ring | 1 | 86 | Male | Caucasian | AMD1 | 3:36 |
| Limbal ring | 2 | 76 | female | Caucasian | AMD1 | 4:17 |
| Limbal ring | 3 | 72 | Male | Caucasian | AMD1 | 3:50 |
| Limbal ring | 4 | 73 | Male | Caucasian | AMD1 | 5:36 |
| Limbal ring | 5 | 73 | Male | Caucasian | AMD1 | 3:49 |
| Neural retina | 1 | 86 | Male | Caucasian | AMD1 | 3:36 |
| Neural retina | 2 | 73 | female | Hispanic | AMD1 | 6:05 |
| Optic nerve | 1 | 86 | Male | Caucasian | AMD1 | 3:36 |
| Optic nerve | 2 | 73 | Male | Caucasian | AMD1 | 5:09 |
| Optic nerve | 3 | 91 | female | Caucasian | AMD2 | 4:21 |
| Optic nerve head | 1 | 86 | Male | Caucasian | AMD1 | 3:36 |
| Optic nerve head | 2 | 82 | Male | Caucasian | AMD1 | 3:34 |
| Optic nerve head | 3 | 78 | Male | Caucasian | AMD1 | 5:43 |

**Supplemental Table 2.** human whole globe donor information (*Angptl7* RNAscope – chromogenic staining)

| Tissue | Donor | Status | Glaucoma grade | Age | Sex | Race | AMD | Death-to-preservation time (hours:minutes) |
| --- | --- | --- | --- | --- | --- | --- | --- | --- |
| Outflow | 1 | Non-glaucoma | N/A | 85 | Male | Caucasian | AMD1 | 4:46 |
| Outflow | 2 | Non-glaucoma | N/A | 78 | Male | Caucasian | AMD1 | 5:02 |
| Outflow | 3 | Non-glaucoma | N/A | 83 | Female | Caucasian | AMD1 | 3:17 |
| Outflow | 4 | Non-glaucoma | N/A | 84 | Male | Caucasian | AMD1 | 4:27 |
| Outflow | 1 | Glaucoma | Severe | 65 | Female | Caucasian | AMD1 | 3:48 |
| Outflow | 2 | Glaucoma | Severe | 97 | Male | Caucasian | AMD3 | 5:30 |
| Outflow | 3 | Glaucoma | Mild | 68 | Female | Caucasian | AMD2 | 5:12 |
| Outflow | 4 | Glaucoma | N/A | 89 | Male | Caucasian | AMD1 | 5:10 |
| Outflow | 5 | Glaucoma | N/A | 91 | Male | Caucasian | AMD1 | 4:24 |
| ONH | 1 | Non-glaucoma | N/A | 85 | Male | Caucasian | AMD1 | 4:46 |
| ONH | 2 | Non-glaucoma | N/A | 78 | Male | Caucasian | AMD1 | 5:02 |
| ONH | 3 | Non-glaucoma | N/A | 77 | Male | Caucasian | AMD2 | 4:06 |
| ONH | 4 | Non-glaucoma | N/A | 77 | Male | Caucasian | AMD1 | 4:05 |
| ONH | 1 | Glaucoma | Mild | 86 | Male | Caucasian | AMD2 | 2:16 |
| ONH | 2 | Glaucoma | N/A | 92 | Female | Caucasian | AMD1 | 3:25 |
| ONH | 3 | Glaucoma | Severe | 76 | Female | Caucasian | AMD2 | 4:50 |
| ONH | 4 | Glaucoma | Mild | 77 | Male | Caucasian | AMD1 | 6:29 |
| ONH | 5 | Glaucoma | Mild | 68 | Female | Caucasian | AMD2 | 5:12 |

**Supplemental Table 3.**
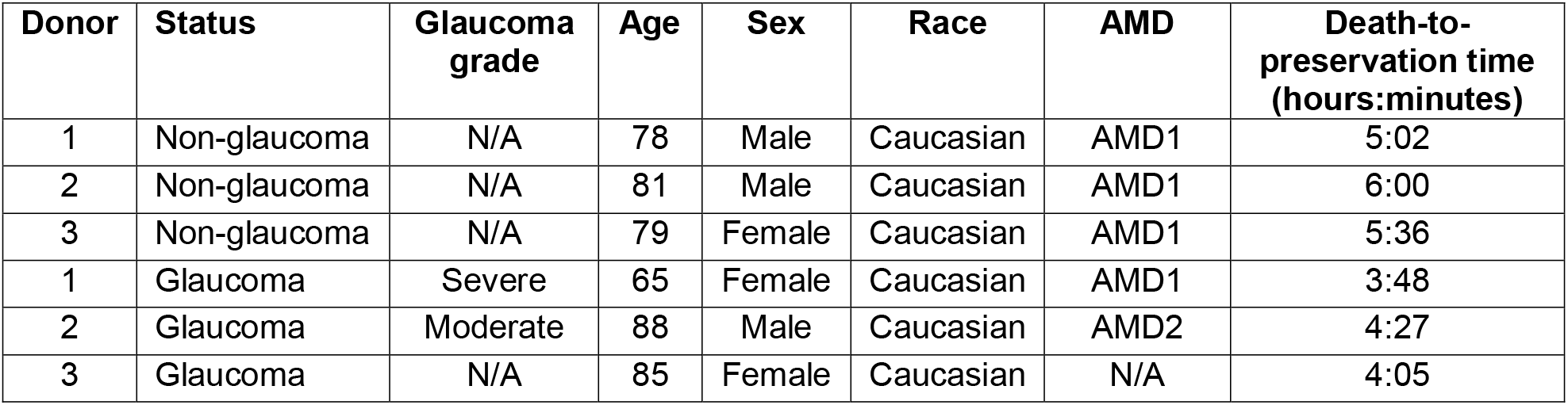
human whole globe donor information (*ANGPTL7*-*GFAP*-IBA1 fluorescent co-staining)

| Donor | Status | Glaucoma grade | Age | Sex | Race | AMD | Death-to-preservation time (hours:minutes) |
| --- | --- | --- | --- | --- | --- | --- | --- |
| 1 | Non-glaucoma | N/A | 78 | Male | Caucasian | AMD1 | 5:02 |
| 2 | Non-glaucoma | N/A | 81 | Male | Caucasian | AMD1 | 6:00 |
| 3 | Non-glaucoma | N/A | 79 | Female | Caucasian | AMD1 | 5:36 |
| 1 | Glaucoma | Severe | 65 | Female | Caucasian | AMD1 | 3:48 |
| 2 | Glaucoma | Moderate | 88 | Male | Caucasian | AMD2 | 4:27 |
| 3 | Glaucoma | N/A | 85 | Female | Caucasian | N/A | 4:05 |

**Supplemental Table 4.** human whole globe donor information (*LAIR1* RNAscope)

| Donor | Status | Glaucoma grade | Age | Sex | Race | AMD | Death-to-preservation time (hours:minutes) |
| --- | --- | --- | --- | --- | --- | --- | --- |
| 1 | Non-glaucoma | N/A | 77 | Male | Caucasian | AMD1 | 4:05 |
| 2 | Non-glaucoma | N/A | 94 | Male | Caucasian | AMD1 | 4:30 |
| 3 | Non-glaucoma | N/A | 83 | Female | Caucasian | AMD1 | 3:17 |
| 4 | Non-glaucoma | N/A | 78 | Male | Caucasian | AMD1 | 5:02 |
| 1 | Glaucoma | Mild | 68 | Female | Caucasian | AMD2 | 5:12 |
| 2 | Glaucoma | N/A | 88 | Male | Caucasian | AMD2 | 4:27 |
| 3 | Glaucoma | N/A | 76 | Female | Caucasian | AMD2 | 4:50 |
| 4 | Glaucoma | Mild | 89 | Male | Caucasian | AMD1 | 5:10 |
| 5 | Glaucoma | N/A | 86 | Male | Caucasian | AMD2 | 2:16 |
| 6 | Glaucoma | N/A | 93 | Male | Caucasian | Dry AMD | 5:12 |
| 7 | Glaucoma | Mild | 77 | Male | Caucasian | AMD1 | 6:29 |

**Supplemental Table 5.** Primary antibodies used for mouse flat-mount imaging.

| Study | Marker | Product | Species | Dilution |
| --- | --- | --- | --- | --- |
| Baseline characterization | BRN3A | Millipore MAB1585 | Mouse | 1:200 |
| Baseline characterization | GFAP | Abcam Ab53554 | Goat | 1:200 |
| Baseline characterization | IBA1 | Wako 019-19741 | Rabbit | 1:1,500 |
| HAMA neuroprotection | BRN3A | Synaptic Systems 411 004 | Guinea pig | 1:2,000 |
| HAMA neuroprotection | GFAP | Invitrogen PA1-10004 | Chicken | 1:4,000 |
| HAMA neuroprotection | IBA1 | Wako 019-19741 | Rabbit | 1:1,500 |

**Supplemental Table 6.** Secondary antibodies used for mouse flat-mount imaging.

| Study | Item | Product | Dilution |
| --- | --- | --- | --- |
| Baseline characterization | Donkey anti-mouse AlexaFluor568 | Thermo Scientific A10037 | 1:500 |
| Baseline characterization | Donkey anti-goat AlexaFluor647 | Thermo Scientific A21447 | 1:500 |
| Baseline characterization | Donkey anti-rabbit AlexaFluor488 | Thermo Scientific A21206 | 1:500 |
| HAMA neuroprotection | Goat anti-guinea pig AlexaFluor568 | Thermo Scientific A11075 | 1:500 |
| HAMA neuroprotection | Goat anti-chicken AlexaFluor647 | Thermo Scientific A21449 | 1:500 |
| HAMA neuroprotection | Donkey anti-rabbit AlexaFluor488 | Thermo Scientific A21206 | 1:500 |

**Supplemental Table 7.** TaqMan probes (qPCR)

| Species | Gene | Reporter | Assay |
| --- | --- | --- | --- |
| Mouse | *Angptl7* | FAM | Mm01256627_g1 |
| Mouse | *Gfap* | FAM | Mm01253033_m1 |
| Mouse | *Krt12* | FAM | Mm07296877_m1 |
| Mouse | *Nrl* | FAM | Mm00476550_m1 |
| Mouse | *Gapdh* | VIC | Mm99999915_g1 |
| Human | *IL1B* | FAM | Hs01555410_m1 |
| Human | *IL1RN* | FAM | Hs00893626_m1 |
| Human | *LAIR1* | FAM | Hs00253790_m1 |
| Human | *LILRB2* | FAM | Hs01629548_s1 |
| Human | *TNF* | FAM | Hs00174128_m1 |
| Human | *GAPDH* | VIC | Hs02786624_g1 |

**Supplemental Table 8.**
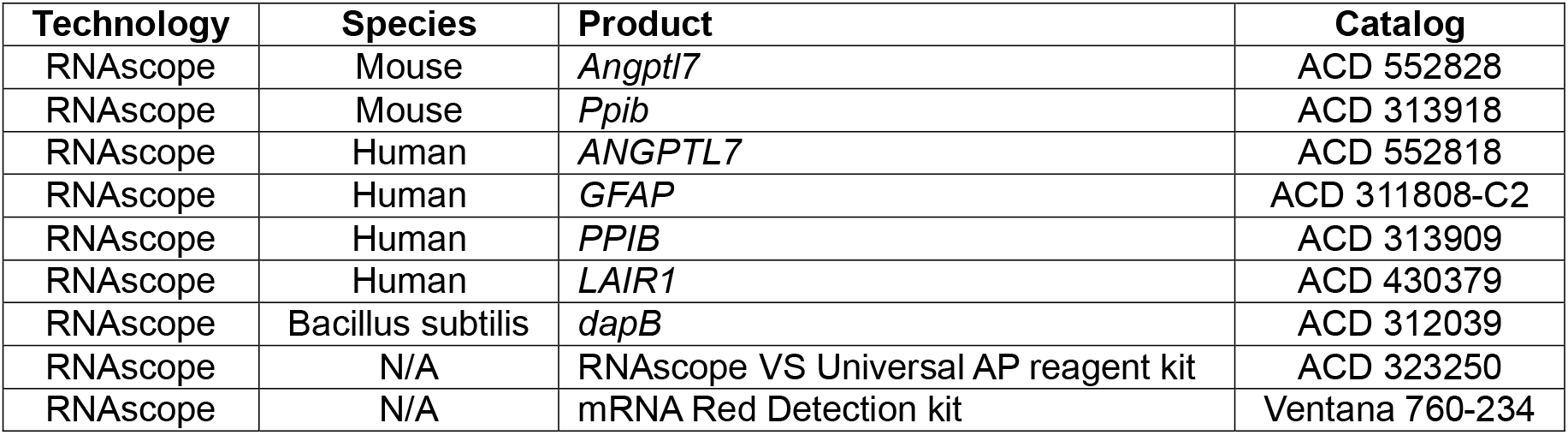
RNAscope reagents.

| Technology | Species | Product | Catalog |
| --- | --- | --- | --- |
| RNAscope | Mouse | *Angptl7* | ACD 552828 |
| RNAscope | Mouse | *Ppib* | ACD 313918 |
| RNAscope | Human | *ANGPTL7* | ACD 552818 |
| RNAscope | Human | *GFAP* | ACD 311808-C2 |
| RNAscope | Human | *PPIB* | ACD 313909 |
| RNAscope | Human | *LAIR1* | ACD 430379 |
| RNAscope | Bacillus subtilis | *dapB* | ACD 312039 |
| RNAscope | N/A | RNAscope VS Universal AP reagent kit | ACD 323250 |
| RNAscope | N/A | mRNA Red Detection kit | Ventana 760-234 |

**Supplemental Table 9.** Additional reagent details.

| Name | Catalog |
| --- | --- |
| Recombinant human ANGPTL7 (6X His tag) | R&D 914-AN |
| Recombinant human ANGPTL2, carrier-free | R&D 9746-AN/CF |
| Recombinant human ANG2 | R&D 623-AN |
| His tag antibody | R&D MAB050 |
| Direct-zol RNA microprep | Zymo R2063 |
| SuperScript™ IV VILO™ Master Mix | Invitrogen 11766050 |
| DNeasy Blood & Tissue kit | Qiagen 69504 |
| Luminex Performance Human XL Cytokine Panel | R&D FCSTM18B |
| Goat Anti-Mouse IgG Antibody, Fc, FITC conjugate | Sigma AP127F |
| Anti-human ANG2 antibody | R&D AF623 |
| Anti-human ANGPTL7 antibody | R&D AF914 |

